# Broad and Durable Humoral Responses Following Single Hydrogel Immunization of SARS-CoV-2 Subunit Vaccine

**DOI:** 10.1101/2022.12.12.520166

**Authors:** Ben S. Ou, Olivia M. Saouaf, Jerry Yan, Theodora U.J. Bruun, Julie Baillet, Xueting Zhou, Neil P. King, Eric A. Appel

## Abstract

Most vaccines require several immunizations to induce robust immunity, and indeed, most SARS-CoV-2 vaccines require an initial two-shot regimen followed by several boosters to maintain efficacy. Such a complex series of immunizations unfortunately increases the cost and complexity of populations-scale vaccination and reduces overall compliance and vaccination rate. In a rapidly evolving pandemic affected by the spread of immune-escaping variants, there is an urgent need to develop vaccines capable of providing robust and durable immunity. In this work, we developed a single immunization SARS-CoV-2 subunit vaccine that could rapidly generate potent, broad, and durable humoral immunity. We leveraged injectable polymer-nanoparticle (PNP) hydrogels as a depot technology for the sustained delivery of a nanoparticle COVID antigen displaying multiple copies of the SARS-CoV-2 receptor-binding-domain (RBD-NP), and potent adjuvants including CpG and 3M-052. Compared to a clinically relevant prime-boost regimen with soluble vaccines formulated with CpG/Alum or 3M-052/Alum adjuvants, PNP hydrogel vaccines more rapidly generated higher, broader, and more durable antibody responses. Additionally, these single-immunization hydrogel-based vaccines elicited potent and consistent neutralizing responses. Overall, we show that PNP hydrogels elicit improved anti-COVID immune responses with only a single administration, demonstrating their potential as critical technologies to enhance our overall pandemic readiness.

## 1. Introduction

The emergence of the SARS-CoV-2 virus, leading to the COVID-19 pandemic, has claimed over 6.5 million deaths globally so far.[1] In only one year, unprecedented scientific achievements resulted in the development and FDA approval of several COVID-19 vaccines. While this striking effort highlights the feat of modern vaccinology, global vaccine inequity remains one of the greatest challenges as low income countries still face difficulties in completing the recommended vaccination schedule.[2] Furthermore, the rise of immune evasive variants of concern, such as the Omicron family of variants (e.g., BA.1, B.1.1.529; BA.2, B.1.1.529.2; BQ.1, B.1.1.529.5.3.1.1.1.1.1), coupled with waning immunity has resulted in partial or limited protection.[3] Consequently, healthy adults in the United States are, at the time of writing, suggested to receive up to two booster shots, including a bivalent booster against the Omicron BA.4/5 (B.1.1.529.4/B.1.1.529.5) variant.[4]

There is, therefore, a crucial need to develop vaccines allowing for better global access while providing durable and broad protection against immune evasive variants. Newly developed DNA and mRNA-based vaccines suffer from high costs, limited manufacturing, and limited resource availabilities, as well as stringent storage conditions, thus limiting their use in low-resource settings. Protein-based subunit vaccines have proven to be successful candidates due to their low manufacturing costs and ability for worldwide distribution as they are more stable and less reliant on the cold chain.[5,6] Thus, subunit vaccines represent an invaluable resource to reduce the gap in vaccine equity. The receptor-binding domain (RBD) of the spike protein on the surface of SARS-CoV-2 has been shown to be an attractive vaccine antigen as it is the epitope responsible for complexing with the host cells to initiate infection. Recently, a prominent multivalent RBD-functional nanoparticle (RBD-NP) antigen utilizing self-assembling protein immunogen, RBD-16GS-I53-50, demonstrated broad and potent binding and neutralizing antibody responses.[3,7–9] This immunogen, alongside AS03 adjuvant, was approved by the Korean Ministry of Food and Drug Safety for inoculating adults 18 years and older.[10]

In addition to the use of more stable subunit vaccine formulations, decreasing the need for booster shots and even potentially reducing the number of shots during the primary immunization series, could be a promising approach. Sustained delivery of antigen(s) has been shown to generate significantly better humoral responses such as increased antibody titers and neutralizing activities. Indeed, Crotty et al. recently demonstrated that sustained HIV Env protein immunogen priming for over two weeks elicited germinal center reactions that last for at least 6 months in non-human primates.[11] In this regard, sustained delivery of SARS-CoV-2 antigens during vaccine priming could eliminate the need for additional boosters. We have previously developed an injectable polymer-nanoparticle (PNP) hydrogel system that enables sustained co-delivery of diverse vaccine cargo upwards of four weeks,[12–15] mimicking natural infections which can result in persisting viral materials in the lymph nodes for several weeks.[16–19] These delivery carriers are inexpensive, scalable, and easy to manufacture, therefore representing ideal single immunization vaccine delivery carriers.

Here, we developed PNP hydrogels containing RBD-NPs to achieve single immunization COVID-19 vaccines. We first characterized and modulated the hydrogels’ mechanical properties to tune sustained antigen delivery kinetics. We then assessed the immunogenicity of PNP hydrogels containing RBD-NPs and different clinically relevant molecular adjuvants, TLR1/2 agonist Pam3CSK4, TLR7/8 agonist 3M-052, TLR9 agonist CpG, and STING agonist cGAMP. We compared single immunization of PNP hydrogels formulated with the best performing groups comprising CpG and 3M-052, with prime-boost soluble vaccines formulated with clinically relevant adjuvant Alum and either CpG or 3M-052. Hydrogel vaccines led to higher and more durable anti-RBD titers and greater breadth of antibody responses against variants of concern. Moreover, sera from hydrogel vaccinated mice showed improved neutralizing ability compared to soluble controls. Together, these encouraging results suggest that PNP hydrogels are promising candidates as single immunization vaccine carriers inducing potent, durable, and broad humoral immune responses.

## 2. Results

### 2.1 Hydrogels for Sustained Vaccine Exposure

We have previously described that PNP hydrogels can improve humoral responses when used as vaccine delivery carriers by sustaining and co-delivering vaccine antigens and adjuvants such as ovalbumin, flu hemagglutinin, and SARS-CoV-2 RBD monomer.[12-14,20,21] These injectable hydrogels are rapidly formed by mixing aqueous solutions of hydrophobically-modified hydroxypropylmethylcellulose (HPMC-C_12_) and biodegradable polymeric nanoparticles (NPs) made of poly(ethylene glycol)-*b*-poly(lactic acid) (PEG-*b*-PLA). Unlike previously reported studies,[12,13] we sought to incorporate clinically used potent nanoparticle antigen to demonstrate PNP hydrogels’ clinical feasibility. In this study, we employed PNP hydrogels to encapsulate the SARS-CoV-2 RBD-NP, RBD-I53-50,[9] to achieve single immunization of a SARS-CoV-2 vaccine by tuning the hydrogel’s viscoelastic mechanical properties as well as screening clinically relevant adjuvants (Figure 1, Table S1). The resulting hydrogel-based vaccines are shear-thinning and self-healing, leading to their facile injection using a standard needle and syringe, followed by the formation of a robust subcutaneous depot. This depot provides sustainable co-release of vaccine cargoes over the course of 2-4 weeks and serves as a local inflammatory niche for immune cell infiltration to enhance antigen processing.[12,14] Cryo-electron microscopy images show the PEG-*b*-PLA nanoparticles evenly distributed in the polymer matrix thereby creating empty spaces for uniform drug trapping (Figure S1).

**Figure 1.**
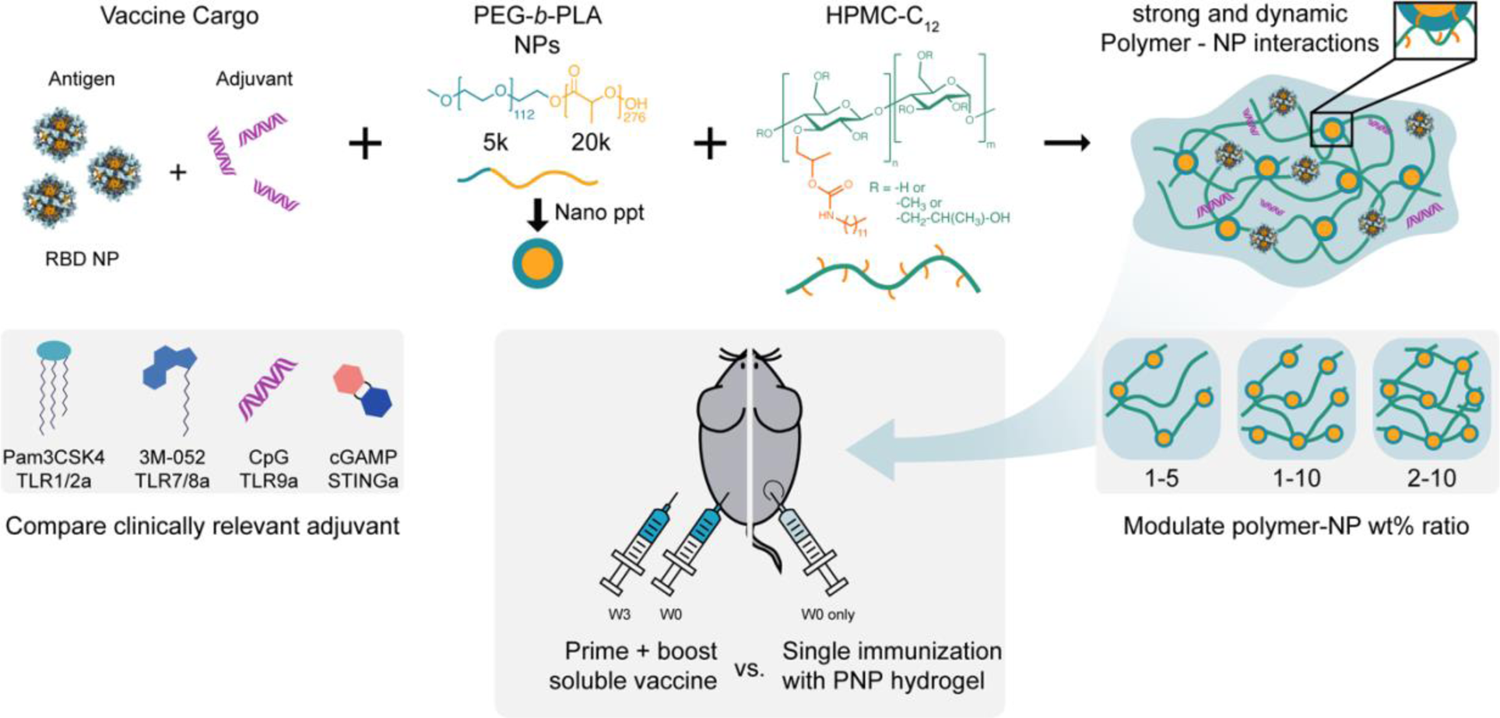
Slow delivery of nanoparticle antigens displaying the SARS-CoV-2 receptor binding domain (RBD-NP) and molecular adjuvants with an injectable depot technology enables potent, broad, and durable COVID immunity. Schematic of injectable polymer-nanoparticle (PNP) hydrogel vaccines where dodecyl-modified hydroxypropylmethylcellulose (HPMC-C_12_) is combined with poly(ethylene glycol)-*b*-poly(lactic acid) nanoparticles (PEG-*b*-PLA NPs) and vaccine cargoes (RBD-NP and clinically relevant molecular adjuvants). Dynamic, multivalent non-covalent interactions between the polymer and the NPs lead to physically crosslinked hydrogels whose unique hierarchical structure enables co-delivery of the vaccine components over user-defined timeframes. The ratio of the polymer to nanoparticles can be tuned to modulate the hydrogel mechanical properties for different vaccine cargo release kinetics.

We hypothesized that tuning the hydrogel viscoelastic properties could impact the release timescale of the large RBD-NP antigen cargo and therefore modulate the induced humoral response. We therefore altered the hydrogel’s network density by varying the weight percent ratio of HPMC-C_12_ to PEG-*b*-PLA NP during mixing (Figure 1). Hydrogel formulations are denoted PNP-X-Y, where X refers to the wt% HPMC-C_12_ and Y refers to the wt% PEG-*b*-PLA NP (*n.b.* the remaining mass is buffer comprising vaccine cargoes). Therefore, a formulation comprising 2 wt% HPMC-C_12_ and 10 wt% PEG-*b*-PLA NP is denoted as PNP-2-10 (*n.b.* the other 88 wt% is buffer).

### 2.2 Shear-Thinning, Self-Healing Hydrogel Characterization

We characterized the viscoelastic properties of PNP-2-10, PNP-1-10, and PNP-1-5 hydrogel formulations using several rheological techniques. We first conducted frequency-dependent oscillatory shear experiments within the linear viscoelastic regime to measure the hydrogels’ frequency responses (Figure 2a). We observed solid-like properties for all formulations, with the storage modulus (G’) being greater than loss modulus (G”) over the range of frequencies tested. Consistent with previous reports,[12,20] we also observed a decrease of both G’ and G” at an angular frequency of ω = 10 rad/s as we decreased the hydrogel network density while Tan(delta) remained unchanged (Figure S2). We then performed flow sweep measurements to test the hydrogels’ shear-thinning behaviors (Figure 2b). We observed a dramatically decreasing viscosity with an increasing shear rate, thereby confirming the hydrogels’ ability to shear-thin, which is necessary for facile injectability. A stress-controlled flow sweep was also performed to measure the hydrogels’ yield stresses, a critical characteristic for maintaining a robust depot upon administration,[12,22] by determining the stress where the viscosity decreases by 3 orders of magnitude (Figure 2c). We observed decreasing yield stresses as network density decreased, suggesting shorter depot lifetimes. Lastly, we conducted step-shear experiments by applying low (0.1 s^-1^) and high (10 s^-1^) shear rates in a stepwise series to assess the hydrogels’ abilities to both shear-thin and self-heal (Figure 2d). When the shear rate returned to the low from the high shear rate, all hydrogels rapidly recovered their mechanical properties and the viscosity increased by two orders of magnitude. This behavior persisted for multiple cycles, confirming the non-covalent crosslinking interactions between the polymer and the NPs. Overall, these studies are consistent with previous findings where different weight precent of HPMC-C_12_ can greatly influence the viscoelastic properties of PNP hydrogels and PEG-*b*-PLA nanoparticles can modulate the stiffness and yield-stresses of the hydrogels.[23] The rheological behavior also confirmed the injectability of all hydrogel formulations that can rapidly self-heal, forming a robust solid-like depot upon injection.

**Figure 2.**
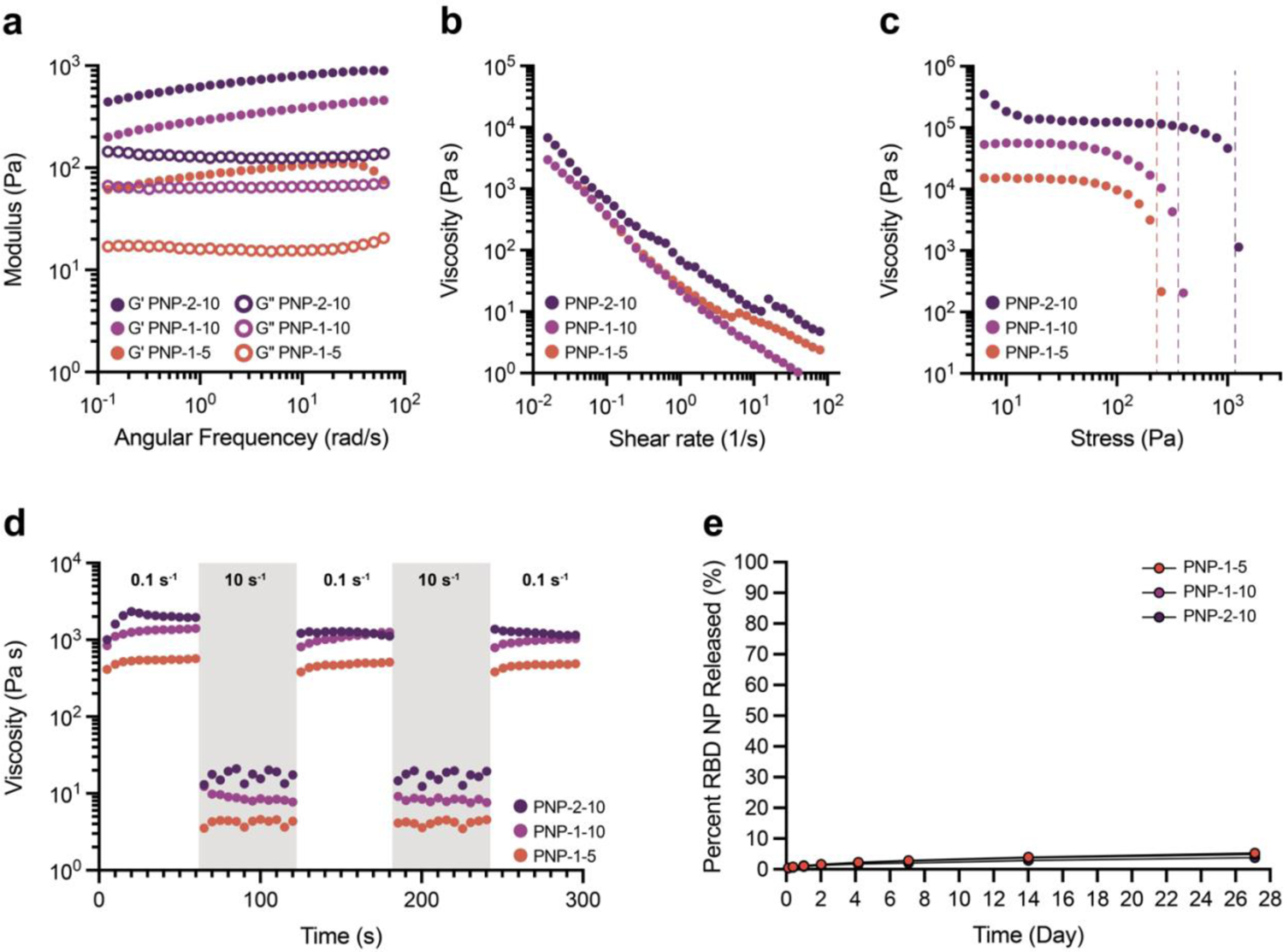
Characterization of Polymer-Nanoparticle (PNP) hydrogels. **(a)** Frequency-dependent oscillatory shear rheology of three different PNP hydrogel formulations. **(b)** Shear-dependent viscosities of PNP hydrogels. **(c)** Stress-controlled oscillatory amplitude sweeps of PNP hydrogels. Yield stresses were determined by the crossover points. **(d)** Step-shear measurements of PNP hydrogels over three cycles of alternating high shear (gray; 10 s^−1^) and low shear (white; 0.1 s^−1^) rates. **(e)** Release kinetics of FITC-Dextran (MW ∼ 500 kDa) of comparable size to RBD-NPs (*D*_H_ ∼ 40nm) from PNP hydrogels in a glass capillary *in vitro* release study.

### 2.3 Kinetics of Cargo Release from the Hydrogels

We have previously determined the mesh size of PNP-2-10 hydrogels to be around 6 nm, and demonstrated that cargo with hydrodynamic radii of similar or larger size to this mesh will be immobilized in the polymer matrix.[21] Therefore, we hypothesized that the RBD-NP antigen, which has a hydrodynamic diameter of 41 nm,[9] would be immobilized by the hydrogel matrix and slowly released upon the erosion of the hydrogel. To test this hypothesis, we determined the *in vitro* release kinetics of 500,000 MW FITC-Dextran (FITC-Dex) as a model cargo, featuring a hydrodynamic diameter close to the antigen (*D*_H_ ∼ 39.2 nm, Table S2). PNP-2-10, PNP-1-10, and PNP-1-5 hydrogels containing FITC-Dex were loaded into capillary tubes and incubated with saline buffer solution at 37 °C. The buffer was completely exchanged at the indicated times for four weeks and the amount of FITC-Dex released into the solution was quantified in each sample (Figure 2e). Less than 6% cumulative release of FITC-Dex was observed for all formulations by week 4, with higher hydrogel network density releasing slightly less cargo. The small percentages of FITC-Dex released suggests that cargo diffusion is limited by the mesh of the hydrogel and that release is primarily by hydrogel erosion in the body, which is severally limited in this *in vitro* setup. *In vivo* erosion of the hydrogels, which can also be measured as the persistence time of the hydrogels in the subcutaneous space, has been previously reported.[22] Half-lives of hydrogel retention were found to be 8.5 days for PNP-1-5 and 10.9 days for PNP-1-10. While the half-life of hydrogel retention was not measured for PNP-2-10, prior work has also demonstrated that yield stress and pre-shear viscosity are respectively predictive of depot formation and depot persistence time. As PNP-2-10 hydrogels exhibit higher yield stress and higher pre-shear viscosities than PNP-1-10 hydrogels (Figure 2c), we expect the depot persistence half-life of PNP-2-10 formulations to be approximately 14 days. Overall, these observations indicated that PNP hydrogels can finely tune cargo diffusion and sustainably release RBD-NP antigen for several weeks.

### 2.4 Vaccine Responses of Different Hydrogel Formulations

To achieve the goal of single administration COVID vaccines by sustained vaccine delivery, we immunized C57BL/6 mice (n = 6) with PNP hydrogel vaccines comprising 3 μg of RBD-NP and collected sera over a ten-week period (Figure 3a). We first assessed how the vaccine release kinetics would influence the humoral responses by immunizing mice with 100 μL of three different hydrogel formulations: PNP-1-5, PNP-1-10, and PNP-2-10, all adjuvanted with CpG (20 μg). Because hydrogel erosion is the main mechanism of RBD-NP release, we further hypothesized that we could extend the hydrogel persistence time and thereby the release of vaccine cargoes by increasing the injection volume. We tested this hypothesis by immunizing mice with twice the volume (200 μL) of PNP-1-5 hydrogel, since its lower yield stress could maximize cell infiltration, while keeping antigen and adjuvant doses consistent (this formulation is referred to as PNP-1-5 2X volume). We observed high antigen-specific total IgG titers over the entire ten-week period for all hydrogel formulations (Figure 3b, Figure S3). Notably, with just one injection, we observed seroconversion across all treated animals in all groups. PNP-1-5 produced the highest titers over time compared to the other groups *(p* values in Table S3). PNP-1-5 was therefore selected as formulation of choice for the next steps of the study. Importantly, like previous reports,[12,24] we did not observe foreign body responses or indications of poor tolerability of PNP hydrogels, demonstrating their excellent biocompatibility (Figure S4-S5).

**Figure 3.**
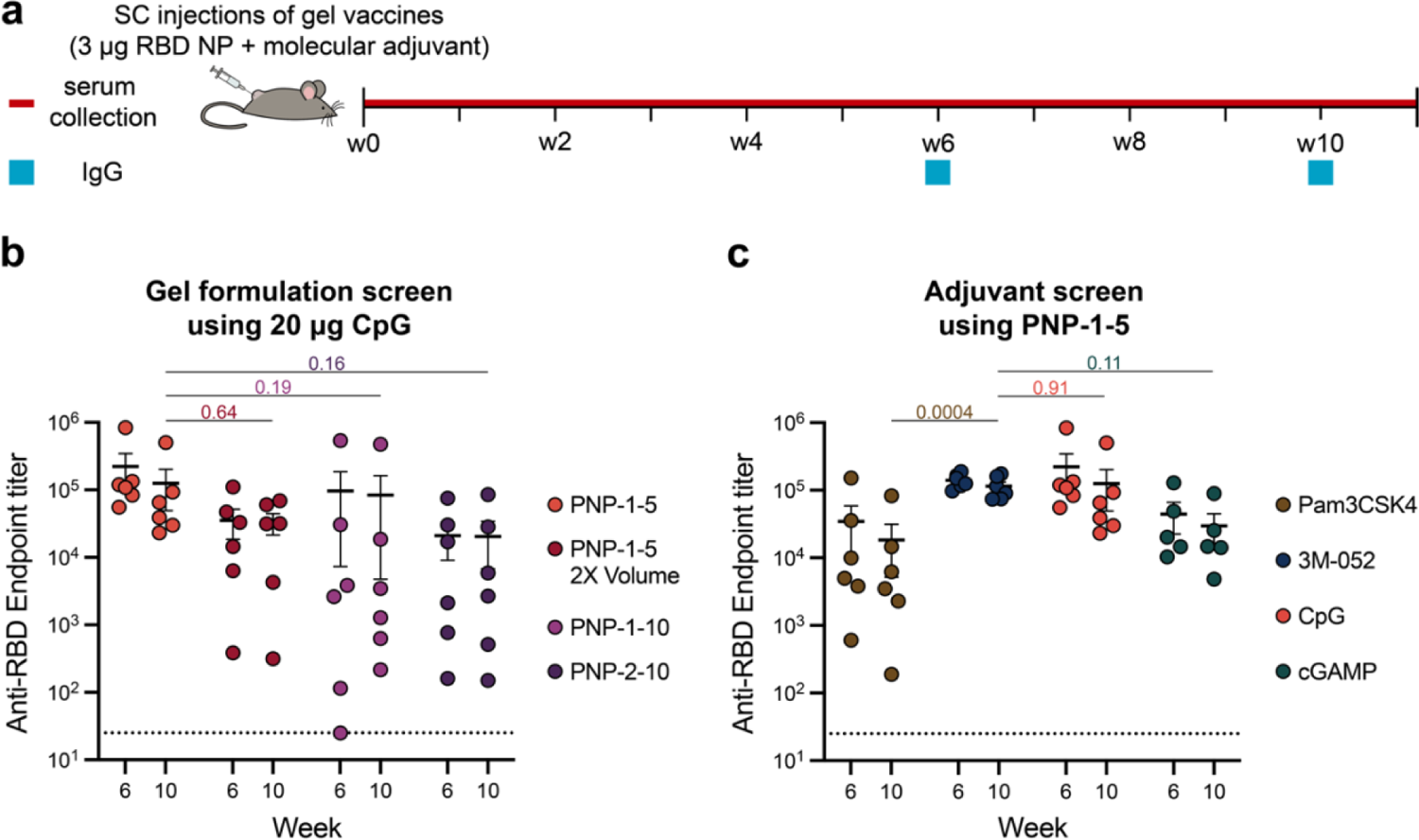
PNP hydrogel vaccines induce robust *in vivo* humoral responses with RBD-NP. **(a)** Timeline of mouse immunizations and blood collection. Mice were immunized with PNP hydrogels formulated with 3 μg of RBD-NP on day 0 and serum was collected over time. **(b)** Anti-RBD IgG binding endpoint titers of different PNP hydrogel formulation vaccinations, all containing 20 μg of CpG. **(c)** Anti-RBD IgG binding endpoint titers of PNP-1-5 formulated with different clinically relevant molecular adjuvants. Data are shown as mean +/- SEM. *p* values were determined using a 2way ANOVA with Tukey’s multiple comparisons test on the logged titer values for IgG titer comparisons. Complete *p* values for comparisons are shown in Tables S3 & S4.

We then evaluated whether utilizing different adjuvants would affect the immune response. We immunized mice (n = 5-6) with PNP-1-5 hydrogels formulated with 3 μg of RBD-NP and one of the four clinically relevant molecular adjuvants: Pam3CSK4 (20 μg), 3M-052 (1 μg), CpG (20 μg), or cGAMP (20 μg) (Figure 3a). Consistent with previous findings,[13] all hydrogel vaccines regardless of adjuvants resulted in seroconversion in all mice with high antibody titers throughout the ten-week study period (Figure 3c, Figure S6). Mice immunized with CpG and 3M-052 adjuvants generated the highest titers, with anti-RBD endpoint titers of 2.2 x 10^5^ and 1.4 x 10^5^ measured on week 6, respectively. Both groups generated more than double the titers measured from the cGAMP group, with an endpoint titer of 4.5 x 10^4^, and significantly higher titers than the Pam3CSK4 group, with an endpoint titer of 3.5 x 10^4^ *(p* values for all comparisons are reported in Table S4). Furthermore, RBD-NP based PNP hydrogel vaccines improved titers durability from Week 6 to Week 10 compared to a RBD monomer hydrogel vaccine previously reported (44% and 78% decrease in titers, respectively; Figure S7; *p* values reported in Table S5).[13] Therefore, RBD-NP hydrogels led to a 3.7-fold higher endpoint titer compared to the RBD monomer hydrogels on Week 10 (*p* values in Table S6). Overall, these studies demonstrated that sustained release of RBD-NP with 3M-052 and CpG in PNP-1-5 hydrogels could maintain robust antibody titers over a period of 10 weeks, thereby mitigating the need for boosting.

We also assessed *in vitro* capillary release of CpG and 3M-052 from the PNP hydrogels. Like the FITC-Dextran *in vitro* release study, we loaded PNP-2-10, PNP-1-10, and PNP-1-5 hydrogels containing CpG and 3M-052 into capillary tubes and incubated them with saline buffer solution at 37 °C. The buffer was completely exchanged at the indicated times for four weeks and the amount of CpG and 3M-052 released into the solution was quantified in each sample (Figure S8a-b). Only around 30% of CpG and 3M-052 was released from the PNP hydrogels over this timeframe, regardless of formulations, suggesting severely limited CpG and 3M-052 diffusion. Notably, these results highlight the ability of the PNP hydrogels to precisely co-deliver vaccine antigens and adjuvants over prolonged timeframes despite their distinct physical and chemical properties.

### 2.5 Single Immunization of Hydrogel Vaccines

We next compared the best performing single-administration RBD-NP PNP hydrogel vaccines with standard prime-boost soluble RBD-NP vaccines in saline solution. We prepared soluble vaccines comprising RBD-NP (1.5 μg), Alum (100 μg), and either CpG (20 μg) or 3M-052 (1 μg), which are referred as Soluble CpG/Alum and Soluble 3M-052/Alum, respectively. Alum was included in the soluble groups as a depot vehicle to better mimic clinical adjuvant formulations and previous studies.[7,8] Soluble vaccines were subcutaneously administrated to C57BL/6 mice (n = 5) at Week 0 (prime) and Week 3 (boost). In this way, all groups received the same total dose of RBD-NP in the immunization schedule (one 3ug hydrogel immunization vs. two 1.5ug soluble immunizations). Sera were collected from Week 1 to Week 10 and compared with mice immunized with PNP-1-5 containing CpG or 3M-052, referred as Gel CpG and Gel 3M-052, respectively (Figure 4a).

**Figure 4.**
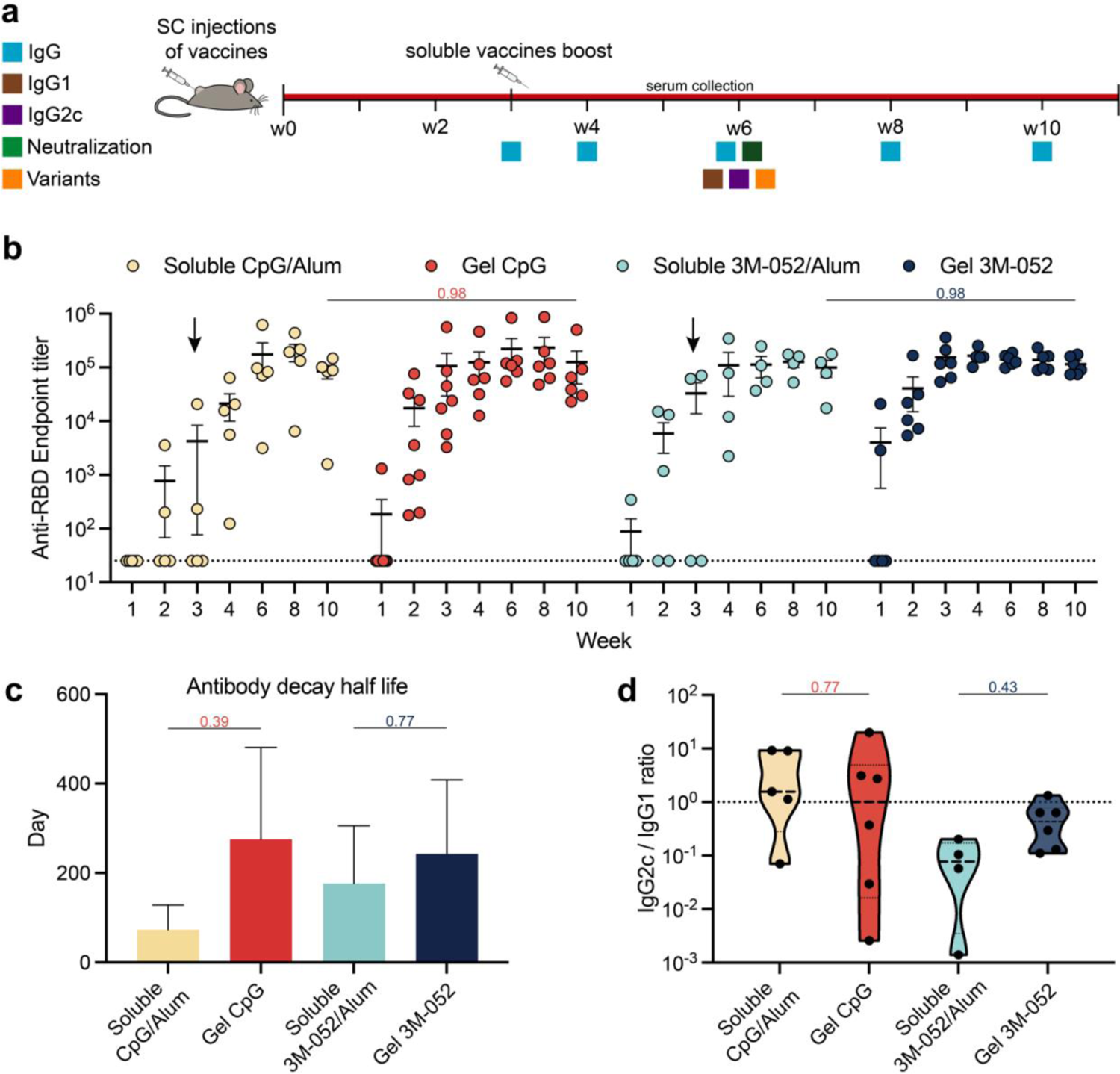
Single immunization hydrogel vaccines elicited comparable antibody titers compared to soluble prime + boosted vaccines. **(a)** Timeline of mouse immunizations and blood collection to determine IgG titers. Mice were immunized with either PNP-1-5 hydrogels formulated with 3 μg of RBD-NP on day 0 or soluble vaccines formulated with 1.5 μg of RBD on day 0 and day 21. IgG1, IgG2c, neutralization, and variants titers were determined on day 42. **(b)** Anti-RBD IgG binding endpoint titers of single immunization PNP hydrogel vaccines formulated with either CpG or 3M-052 alongside soluble vaccine controls before and after boosting (indicated by an arrow). **(c)** A power law decay model was used to determine the decay half-lives of the binding antibodies for the treatment groups over time. **(d)** The ratio of Anti-RBD IgG2c to IgG1 titers from serum collected on week 6 after first immunization. Lower values suggest a Th2-skewed humoral response, whereas higher values suggest a Th1-skewed cellular response. Data are shown as mean +/- SEM. *p* values were determined using a 2way ANOVA with Tukey’s multiple comparisons test on the logged titer values for IgG titer comparisons. Complete *p* values for comparisons are shown in Tables S7-S12.

We first measured anti-RBD total IgG titers over time (Figure 4b). On Week 1, several mice immunized with hydrogel vaccines (Gel CpG and Gel 3M-052) had already seroconverted and by Week 2, all mice in the hydrogel groups had detectable antibody titers. Mice vaccinated with hydrogel vaccines elicited antibody endpoint titers averaging at 1.4 x 10^5^ and 1.5 x 10^5^, respectively on Week 3 (*p* values in Table S7). In contrast, even on Week 3, soluble vaccines (Soluble CpG/Alum and Soluble 3M-052/Alum) produced highly varied antibody titers with more than half the groups’ titers below the limit of detection. Moreover, PNP hydrogel vaccines maintained higher titers at all time points when compared to soluble vaccines, even after boosting. For example, on Week 6, the average endpoint titer for Gel CpG was 2.2 x 10^5^, while it was 1.7 x 10^5^ for Soluble CpG/Alum. Similar trends were observed for Gel 3M-052 versus Soluble 3M-052/Alum. We then evaluated the duration of immunity by collecting the sera 6 months after initial immunization of Soluble CpG/Alum and Gel CpG. Notably, Gel CpG maintained 56.5% of antibody titers from Week 10 which was higher than Soluble CpG/Alum, only retaining 21.7% (Figure S9a-b, *p* values in Table S8). Notably, the average endpoint titer of Gel CpG at 6 months post-immunization, 1.1 x 10^5^, was higher than the early endpoint titer of Soluble CpG at Week 10. Interestingly, Soluble and Gel 3M-052 groups maintained similar binding titers at 6 months post-immunization (Figure S9b, *p* values in Table S8). We estimated the half-life of binding antibody titers using a power law decay model, which has been shown to accurately reflect the kinetics of antibody response *in vivo*.[3] The estimated half-lives were found to be 67 days and 270 days after the peak of titers (determined to be Day 56) for soluble CpG/Alum and Gel CpG, respectively (Figure 4c, Figure S9c, *p* values are shown in Table S9). In contrast, the estimated half-lives were found to be 177 days and 243 days after the peak for soluble 3M-052/Alum and Gel 3M-052, respectively (Figure 4c, Figure S9d, *p* values are shown in Table S9). While the hydrogel groups maintained similar decay half-lives regardless of the adjuvants, we observed higher decay variability for both soluble vaccine groups and generally longer decay half-lives for soluble 3M-052/Alum than CpG/Alum. These data suggest that the single immunization hydrogel treatments were able to reliably sustain antibody titers over a prolonged period.

We then assessed the IgG isotypes that made up the total IgG titer of each group on Week 6 to determine whether the choice of adjuvants or hydrogels would influence immune signaling. Specifically, we evaluated the titers of IgG1 and IgG2c isotypes as these isotypes are strong indicators of Th2- and Th1-skewed immune responses, respectively.[25] Soluble CpG/Alum had the lowest IgG1 response while Gel CpG had more than 5-fold the titer (Figure S9e, *p* values are shown in Table S10). Conversely, Soluble 3M-052/Alum had the lowest IgG2c endpoint titer while CpG-containing groups elevated IgG2c titers by 6.7-fold for Soluble CpG and 4.4-fold for Gel CpG (Figure S9f, *p* values are shown in Table S11). Elevated IgG2c titers for the CpG groups raised the IgG2c/IgG1 ratio to close to 1, suggesting a balanced Th1/Th2 response, which is consistent with previous findings including CpG as an adjuvant (Figure 4d, *p* values are shown in Table S12).[8,13,26,27] With low IgG2c titer, Soluble 3M-052/Alum has an IgG2c/IgG1 ratio less than 1, suggesting a Th2-skewed response. However, Gel 3M-052 maintained a high IgG2c endpoint titer, leading to a more balanced IgG2c/IgG1 ratio and Th1/Th2 responses. The balanced Th1/Th2 response from the hydrogel groups is ideal as it has been shown to generate a favorable COVID-19 disease outcome.[28,29]

We then measured the anti-spike IgG titers at Week 6 to confirm the antibodies elicited from the RBD-NP vaccines cross-reacted with RBD presented on the native SARS-CoV-2 spike proteins. Further, we evaluated the breadth of protection generated by different vaccine groups by assessing the antibody titers against SARS-CoV-2 variants known to be immune escape variants, including Delta (B.1.617.2) and Omicron (B.1.1.529) (Figure 5, *p* values are shown in Table S13).[30] Across the groups, wild-type spike (WT Spike) titers reflected a similar trend to those observed with anti-RBD endpoint titers. Mainly, Gel CpG and Gel 3M-052 had the highest anti-WT Spike endpoint titers with little variation among the individual mice. On the contrary, higher deviations and lower titers were observed for the soluble vaccine groups. When assessing the endpoint titers against the Delta variant as compared to the WT spike titers, less than 1.5-fold decreases were measured for Soluble CpG/Alum, Gel CpG, and Gel 3M-052, with the hydrogel groups having the highest titers. Interestingly, a 5.9-fold decrease of titers between the Delta variant and the WT was measured for Soluble 3M-052/Alum. Although endpoint titers against the Omicron variant across all vaccine groups decreased significantly when compared to their wildtype spike titers, we observed fewer titer variabilities for the hydrogel groups with Gel 3M-052, which demonstrated the highest anti-Omicron titer. Notably, sera from two mice vaccinated with Soluble CpG/Alum had titers against Omicron below the limit of detection. From these observations, the hydrogel vaccines demonstrated improved breadth and durability of humoral responses in a single immunization, even against novel variants of concern, compared to soluble prime-boost controls.

**Figure 5.**
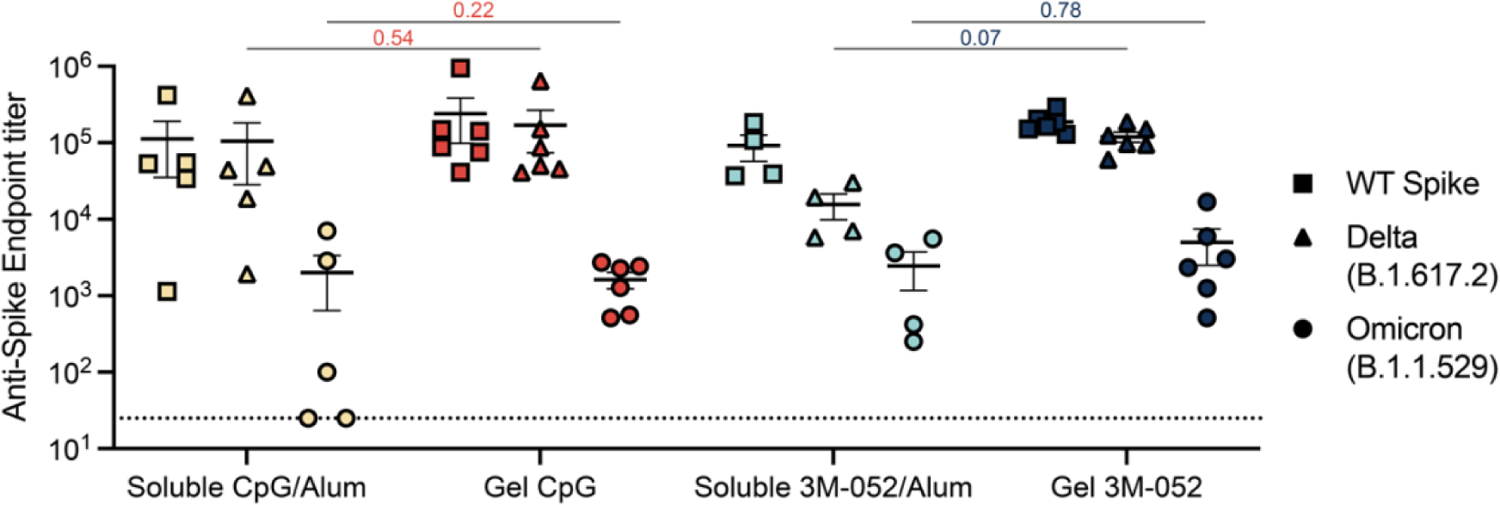
Hydrogel vaccines provide a broader response against SARS-CoV-2 variants of concern. Anti-spike IgG binding endpoint titers from serum collected on week 6 after the initial immunization. Titers were determined for wildtype WT spike as well as Delta (B.1.617.2), and Omicron (B.1.1.529) variants of the spike protein. Data are shown as mean +/- SEM. *p* values were determined using a 2way ANOVA with Tukey’s multiple comparisons test on the logged titer values for IgG titer comparisons. Complete *p* values for comparisons are shown in Table S13.

### 2.6 SARS-CoV-2 Spike-Pseudotyped Viral Neutralization Assay

After confirming that single immunization hydrogel vaccines generated broad and sustained antibody titers, we sought to determine the neutralizing activity of the sera. A lentivirus pseudotyped with SARS-CoV-2 spike was used to measure serum-mediated inhibition of viral entry into HeLa cells overexpressing ACE2 and TMPRSS2.[31,32] We assessed Week 6 serum neutralization by evaluating neutralizing activities of a range of sera concentrations of both soluble and hydrogel vaccines to determine the half-maximal inhibition of infectivity (NT_50_) (Figure 6, *p* values are shown in Table S14). Sera from the Soluble CpG/Alum group presented high variability in neutralizing activities post-boost, with two samples at the lower limit of detection. Conversely, robust neutralizing activities were observed in sera from both Gel CpG and Gel 3M-052 groups, with only one sample in Gel CpG below the FDA’s recommendation for “high titer” classification (NT_50_ ∼ 10^2.4^).[33] We then compared the vaccine groups’ sera neutralizing activities to previously reported human patients’ convalescent sera (Figure S10, *p* values in Table S14).[13] All treatment groups measured higher neutralization titers compared to human patients who were infected with WT COVID-19. We also plotted anti-RBD and anti-Spike binding titers against neutralization NT_50_ titers (Figure S11). We determined a positive correlation between binding titers and neutralization titers (Pearson r = 0.76 for anti-RBD vs. NT_50_ and r = 0.74 for anti-spike vs. NT_50_) further demonstrating the robustness of the binding titers measured using ELISAs.

**Figure 6.**
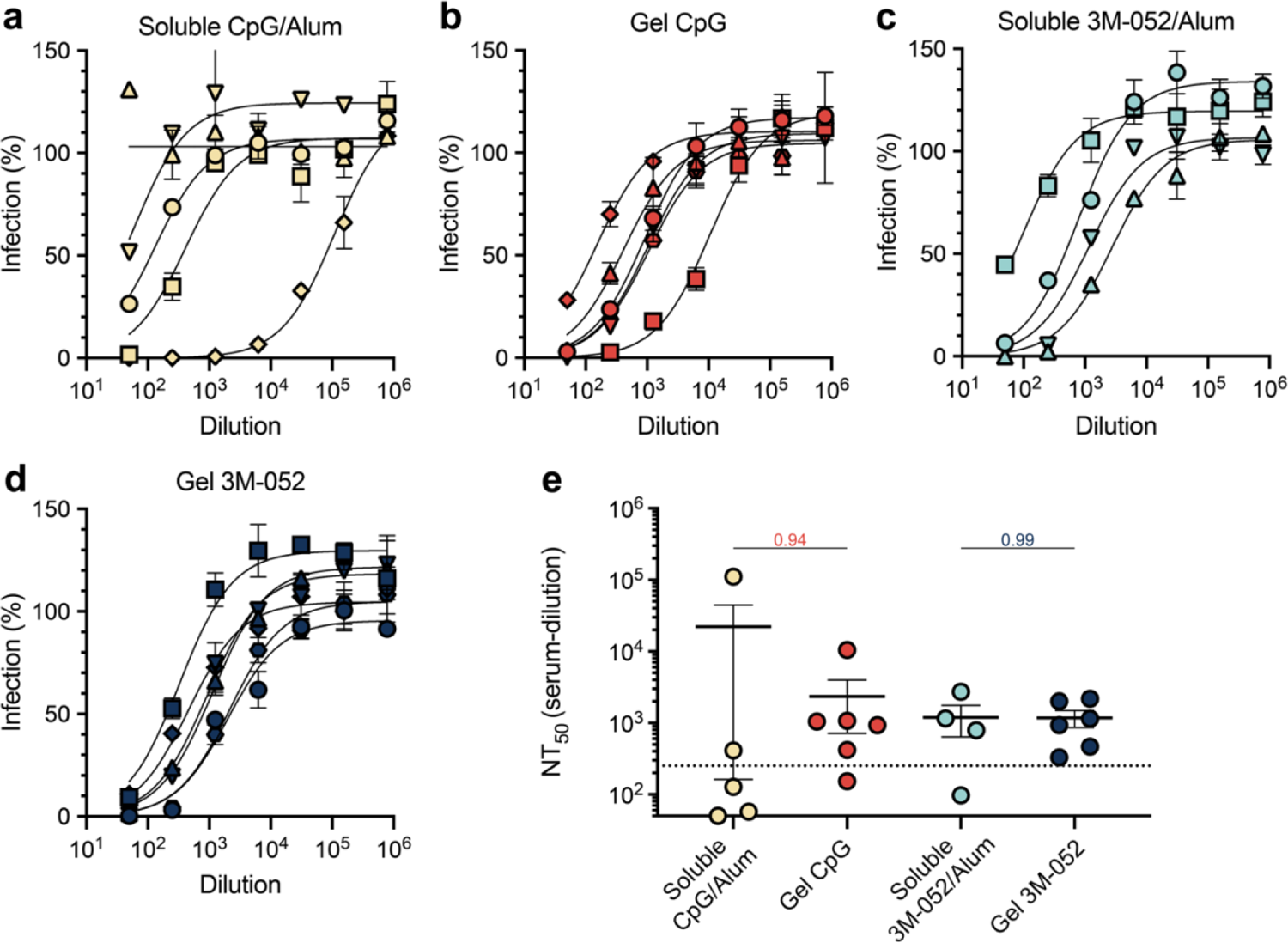
Single immunization hydrogel vaccines elicit robust neutralizing antibodies in mice. **(a-d)** Percent infectivity for all treatment groups at a range of Week 6 serum dilutions as determined by a SARS-CoV-2 spike-pseudotyped viral neutralization assay. **(e)** Comparison of NT_50_ values determined from neutralization curves. Dotted line denote the threshold for which the FDA considers as “high titer”.[33] Data are shown as mean +/- SEM. *p* values were determined using a 2way ANOVA with Tukey’s multiple comparisons test on the logged titer values for IgG titer comparisons. Complete *P* values for comparisons are shown in Table S14.

Taken together, our data suggest that single immunization hydrogel vaccines elicited comparable humoral responses compared to soluble prime-boost RBD-NP vaccines adjuvanted with CpG or 3M-052 and an Alum vehicle. Notably, we observed higher and more durable antibody binding titers with balanced Th1/Th2 responses regardless of the nature of the adjuvants. This led to increased protection against variants of concern and less variable neutralization titers, further demonstrating the potency of PNP hydrogel vaccines.

## 3. Discussion

Waning immunity post-vaccination poses a difficult challenge when managing a rapidly evolving pandemic. At the time of writing, healthy adults in the United States are only considered fully vaccinated after undergoing a prime-boost series of COVID-19 vaccination followed by an additional bivalent booster against the Omicron variant.[4] Unfortunately, only 33% of Americans and 31% of the population worldwide have received at least one booster shot.[34] As a soluble vaccine, RBD-NP adjuvanted with AS03 has been shown to extend antibody titers and provide more durable Omicron variant protection compared to Pfizer-BioNTech and Moderna mRNA vaccines, thus potentially extending the period of protection.[3] Nonetheless, by 6 months, the binding antibody titers decreased to pre-booster magnitude and additional boosters were necessary to confer robust protection.[3] While there are few reports on the development of single-shot nanovaccines,[35–37] an antigen-agnostic approach is necessary for a rapidly employable vaccine platform to combat emerging infectious diseases. Therefore, the present study reports the development of a new vaccine platform strategy that could elicit potent, broad, and durable humoral responses without the need for booster shots. A single injection with the ability to rapidly reach protective levels of neutralizing antibodies is crucial in a pandemic response to protect broad populations more effectively. In this regard, we demonstrated that an injectable PNP hydrogel platform could co-deliver physicochemically distinct vaccine cargo over prolonged timeframes to extend the durability and expand the breadth of immune responses after a single immunization.

We have previously shown that PNP hydrogels improve vaccine humoral responses due to sustained antigen delivery which better mimics antigen exposure during a natural infection.[12–14] We hypothesized here that utilizing this technology for the sustained delivery of a potent SARS-CoV-2 antigen could synergistically improve the durability and breadth of antibody responses from even a single immunization. The RBD-NP used in this study, RBD-16GS-I53-50, has been demonstrated to be stable in 22-27 °C for up to 28 days.[9] Moreover, we have demonstrated previously that nanoparticles can easily be entrapped by the polymer mesh of the PNP hydrogel, allowing for precise control over the kinetics of release by tuning the hydrogel viscoelastic properties. As nanoparticles in the range of 20-200 nm have been shown to increase uptake by APCs and improve lymph node drainage and targeting,[38–50] we speculated that hydrogel formulations providing intermediate rates of erosion (∼2 weeks rather than ∼4 weeks) would allow more RBD-NP to drain selectively to the lymph nodes while synergistically retaining the ability for immune cells to infiltrate the hydrogel depot for antigen processing. In this regard, we found that PNP-1-5 hydrogel formulations produced the highest antibody titers and greatest breadth of humoral responses.

We have also previously demonstrated that PNP hydrogels can deliver many physicochemically distinct molecules over predetermined timeframes, allowing for a screening of clinically relevant adjuvants while controlling for pharmacokinetic effects.[13] Other reports have found that soluble RBD-NP vaccines comprising CpG/Alum or AS03 elicited the strongest immune responses both in mice and NHPs.[7,8] Our findings suggesting that PNP hydrogels formulated with CpG yielded robust immune responses corroborate these previous observations. Using soluble CpG/Alum in our current study as a bridge to compare with previous studies, we can highlight that our hydrogel vaccines may elicit significantly better responses than other clinically relevant adjuvants evaluated previously in the literature, including AS37, Essai O/W 1849101, AddaVax, and Alum alone. Further, like our previous reports evaluating other TLR7/8 agonists with PNP hydrogels,[15,51] we found that 3M-052 – a potent TLR7/8 agonist – elicited robust and durable humoral immunity. In this work, we demonstrated that both CpG and 3M-052 molecular adjuvants are strongly retained within the PNP hydrogels, likely because of non-covalent interactions with the PEG-*b*-PLA NPs, thereby enabling co-delivery of the nanoparticle antigens that are over 100-fold larger and physically embedded within the PNP hydrogel structure. These observations highlight the unique structural characteristics of PNP hydrogels allowing for encapsulation and co-delivery of physicochemically-distinct molecules. We believe that prolonged co-delivery of immune activating adjuvants with antigens can inhibit tolerogenic responses to the antigen. Furthermore, while Alum’s main mechanism is still not fully understood, Alum has been shown to serve as a depot for antigen and adjuvant adsorptions and also induce a strong Th2-skewed response.[52] Since PNP hydrogels already act as an inflammatory niche, we chose not to include Alum within the hydrogel systems to achieve a more balanced Th1 and Th2 response. Soluble vaccine controls, however, included Alum to better mimic clinical formulations.[53–60]

Our present work focuses on humoral immunity because potent neutralizing antibody responses have been found to be strongly correlated with protection against severe COVID-19 disease.[61] Future studies will uncover whether the balanced Th1/Th2 responses in the PNP hydrogels, regardless of the nature of the adjuvant, have any impact on cellular immunity. Such studies could reveal a contrast to the previously reported, predominately Th1-skewed CD4 T cell response found in mice immunized with soluble vaccines comprising RBD-NP and CpG/Alum[8] or the Th2-skewed response we have observed with mice immunized with Soluble 3M-052/Alum.

Crucially, several previous studies have found little to no binding and neutralizing titers against the Omicron BA.1 and BA.2 variants after initial prime-boost immunization series with RBD-NP soluble vaccines.[3,9] Consistent with these findings, we observed several animals in the Soluble CpG/Alum group were non-responders to Omicron variants. In contrast, all hydrogel-based vaccines elicited robust Omicron binding titers. While we have previously reported a preliminary study on single immunization with RBD monomer based hydrogel vaccines, we did not evaluate responses against variants of concern.[62] Future studies will reveal whether our RBD-NP based hydrogel vaccine groups elicit robust neutralizing titers against Omicron and other variants of concern. Several studies have also suggested somatic hypermutation and affinity maturation of memory B cells can persist long after the second immunization (boost) in soluble vaccines.[3] While we did not boost hydrogel vaccines in this study, we have previously shown that PNP hydrogel vaccines improve the magnitude and duration of germinal center reactions and can enhance affinity maturation by upwards of 1000-fold compared to soluble vaccines,[19,63,64] suggesting that future studies into the ultimate extent of somatic hypermutation following a single hydrogel immunization may be of interest. Further investigation could corroborate others’ findings on persistent germinal center B cell activities after sustained HIV immunogen priming.[11]

Supported by the World Health Organization (WHO), the Coalition for Epidemic Preparedness Innovations (CEPI) has financially engaged in the development and manufacturing of vaccines within 100 days in response to “Disease X,” an infectious agent currently unknown to cause human disease.[65,66] We have demonstrated that PNP hydrogels have the unique ability to encapsulate diverse immunogens and molecular adjuvants and enable their sustained co-release to elicit rapid and robust neutralizing immunity in a single administration, reducing both the burden of boosting and the speed to full protection, both of which are features that will be essential in fighting against Disease X. This report is also, to our knowledge, the first to describe sustained delivery technology incorporating clinically used nanoparticle antigens, further demonstrating the clinical feasibility and readiness of PNP hydrogels. The high versatility and robustness offered by PNP hydrogels as a vaccine delivery technology could greatly improve our readiness for “Disease X” threats.

## 4. Conclusion

In this work we report the development of robust candidate single immunization PNP hydrogel COVID vaccines consisting of a SARS-CoV-2 RBD-NP antigen and CpG or 3M-052 adjuvants that elicited potent, broad, and durable humoral immune responses. Compared to dose-matched soluble prime-boost RBD-NP vaccines containing either CpG/Alum or 3M-052/Alum, single administration PNP hydrogel vaccines maintained higher antibody titers over 6 months, elicited more balanced Th1/Th2 responses, generated superior breadth against variants of concern, and produced more consistent neutralizing antibody titers. This robust single immunization strategy could be easily applied to other vaccines for which burdensome boosters limit their worldwide distribution. Further, this slow delivery vaccine delivery platform has the potential to accelerate our readiness for combating Disease X.

## 5. Materials and Methods

### 5.1 Materials

Poly(ethylene glycol)methyl ether 5000 Da (PEG), 3,6-Dimethyl-1,4-dioxane-2,5-dione (Lactide), 1,8-diazabicyclo(5.4.0)undec-7-ene (DBU, 98%), (Hydroxypropyl)methyl cellulose (HPMC, meets USP testing specifications), *N,N*-Diisopropylethylamine (Hunig’s base), *N*-methyl-2-pyrrolidone (NMP), 1-dodecyl isocyanate (99%), mini Quick Spin Oligo columns (Sephadex G-25 Superfine packing material), Sepharose CL-6B crosslinked, bovine serum albumin (BSA), Acetonitrile (ACN), Dimethyl Sulfoxide (DMSO), and dichloromethane (DCM) were purchased from Sigma-Aldrich. RBD-NP was kindly provided by the University of Washington. Pam3CSK4 (Vac-pms), CpG1826 (Vac-1826), 2’3’-cGAMP (Vac-nacga23), and Alum (Alhydrogel 2%; vac-alu) were purchased from Invivogen. 3M-052 and 3M-052/Alum (AAHI-AL030) were purchased from 3M and the Access to Advanced Health Institute (AAHI). SARS-CoV-2 proteins were purchased from Sino Biological including SARS-CoV-2 RBD protein (40592-V08H), SARS-CoV-2 spike protein (40589-V08H4), Alpha B.1.1.7 spike (40591-V08H10), Beta B.1.351 spike (40591-V08H12), Delta B.1.617.2 spike (40591-V08H23), and Omicron B.1.1.529 spike (Sino Biological 40591-V08H41). Goat anti-mouse IgG Fc secondary antibody (A16084) HRP (Horseradish peroxidase) was purchased from Invitrogen. Goat anti-mouse IgG1 and IgG2c Fc secondary antibodies (ab97250, ab97255) HRP were purchased from Abcam. 3,3’,5,5’-Tetramethylbenzidine (TMB) ELISA Substrate, high sensitivity was acquired from Abcam. HIS Lite Cy3 Bis NTA-Ni Complex was purchased from AAT Bioquest. Unless otherwise stated, all chemicals were used as received without further purification.

### 5.2 Preparation of HPMC-C_12_

HPMC-C_12_ was prepared according to a previously reported procedure.[20,67] Briefly, hypromellose (HPMC, 1.5 g) was dissolved in 60 mL of anhydrous NMP. The solution was then heated at 50 °C for 30 mins. A solution of dodecyl isocyanate (0.75 mmol, 183 μL) in 5 mL of anhydrous NMP was added dropwise followed by 105 μL of *N*,*N*-diisopropylethylamine (0.06 mmol). The solution was stirred at room temperature for 20 h. The polymer was recovered from precipitation in acetone and filtered. The polymer was purified through dialysis (3 kDa mesh) in Milli-Q water for 4 days and lyophilized to yield a white amorphous polymer. The polymer mixture was then lyophilized and reconstituted to a 60 mg/mL solution in sterile PBS 1X.

### 5.3 Preparation of PEG-PLA NPs

PEG-*b*-PLA was prepared as previously reported.[20] Prior to use, commercial lactide was recrystallized in ethyl acetate and DCM was dried via cryo distillation. PEG-methyl ether (5 kDa, 0.25 g, 4.1 mmol) and DBU (15 µL, 0.1 mmol) were dissolved in 1 mL of dry DCM under nitrogen atmosphere. Lactide (1.0 g, 6.9 mmol) was dissolved in 4.5 mL of dry DCM under nitrogen atmosphere. The lactide solution was then quickly added to the PEG/DBU mixture and was allowed to polymerize for 8 min at room temperature. The reaction was then quenched with an acetic acid solution and the polymer precipitated into a 1:1 mixture of ethyl ether and hexanes, collected by centrifugation, and dried under vacuum. NMR spectroscopic data, M_n_, and dispersity was then confirmed to match with those previously described.

PEG-*b*-PLA NPs were prepared as previously described.[12,68] A 1 mL solution of PEG-b-PLA in 75:25 ACN:DMSO (50 mg/mL) was added dropwise to 10 mL of Milli-Q water stirring at 600 rpm. The hydrodynamic diameter of the NPs was measured on a DynaPro II plate reader (Wyatt Technology). Three independent measurements were performed for each sample. The particle solution was purified in centrifugal filters (Amicon Ultra, MWCO 10 kDa) at 4500 RCF for 1 h and resuspended in PBS 1X to reach a final concentration of 200 mg/mL.

### 5.4 Preparation of RBD-16GS-I53-5 RBD-NPs

The nanoparticle immunogen (RBD-NP) components and the nanoparticle production method have been previously described.[9] For the purpose of this study, nanoparticles were suspended in the following buffer condition: 50 mM Tris pH 8, 150 mM NaCl, 100 mM l-arginine, and 5% sucrose.

### 5.5 Preparation and Formulations of PNP Hydrogels

Polymer-nanoparticle (PNP) hydrogels were formed at varying (1, 2 wt%) HPMC-C_12_ and (5, 10 wt%) mixture of PEG-*b*-PLA NPs in PBS 1X. Three formulations were tested: (i) PNP-2-10 comprising 2 wt% HPMC-C_12_ and 10 wt% PEG-*b*-PLA NPs, (ii) PNP-1-10 comprising 1 wt% HPMC-C_12_ and 10 wt% PEG-*b*-PLA NPs, and (iii) PNP-1-5 comprising 1 wt% HPMC-C_12_ and 5 wt% PEG-*b*-PLA NPs. Hydrogels were prepared by mixing the corresponding volume of 6 wt% HPMC-C_12_ solution, 20 wt% NPs solution, and TBS 1X to achieve the formulations described above. Based on the desired adjuvant formulations, aqueous adjuvant (Pam3CSK4, 3M-052, CpG1826, or cGAMP) and RBD-NP were included by subtracting the volume of the antigen and adjuvant cargoes from the volume of TBS 1X. The hydrogels were formed by mixing the solutions using syringes connected through an elbow mixer as previously reported.[12]

### 5.6 Hydrogel Rheological Characterization

Rheological characterization was completed on a Discovery HR-2 Rheometer (TA Instruments). Measurements were performed using a 20 mm serrated plate geometry at 25 °C and at 500 µm gap height. Dynamic oscillatory frequency sweeps were conducted at a constant 1% strain and angular frequencies from 0.1 to 100 rad/s. Amplitude sweeps were performed at a constant angular frequency of 10 rad/s from 0.5% to 10,000% strain. Flow sweep and steady shear experiments were performed at shear rates from 50 to 0.005 1/s, whereas stress-controlled flow sweep measurements were conducted at shear rates from 0.001 to 10 1/s. Step-shear experiments were performed by alternating between low shear rates (0.1 rad/s for 60 s) and high shear rates (10 rad/s for 30 s) for three full cycles. Yield stress values were extrapolated from stress-controlled flow sweep and amplitude sweep measurements.

### 5.7 Cargo Release Study

HPMC-C_12_ and PEG-*b-*PLA NPs were mixed with 200 μg of 500,000 MW FITC-Dextran in PBS 1X, CpG in TBS 1X, or 3M-052 in TBS 1X to achieve the hydrogel formulations described above. Glass capillary tubes were plugged at one end with epoxy and 100 μL of hydrogel was injected into the bottom of 4 separate tubes per hydrogel formulation. 400 μL PBS 1X or TBS 1X was added on top of each hydrogel. Tubes were stored upright in an incubator at 37 °C for about 3 weeks. At each time point, ∼400 μL of PBS 1X or TBS 1X was removed and the same amount was replaced. The amount of FITC-Dextran released at each timepoint was determined by measurement of fluorescence with an excitation of 480 nm and an emission of 520 nm. The amount of CpG was measured by absorbance at 260nm and the amount of 3M-052 was measured by absorbance at 230nm. The fluorescence and absorbance measurements were fitted with standard curves of known FITC-Dextran, CpG, or 3M-052 concentrations. The cumulative release was calculated and normalized to the total amount released over the duration of the experiment.

### 5.8 Vaccine Formulations

The vaccines contained 1.5 μg of RBD-NP per dose and varying amounts of adjuvants in either soluble form or in PNP hydrogels. The formulations are outlined in Table S1. Soluble vaccines received one dose of antigen and either a dose of adjuvant of CpG1826/Alum (20 μg + 100 μg, respectively) or 3M-052/Alum (1 μg + 100 μg, respectively). Hydrogel vaccines received two (double) doses of antigen and a dose adjuvant (20 μg of Pam3CSK4, CpG1826, cGAMP, or 1 μg of 3M-052). For soluble groups, vaccines were prepared in TBS 1X to a volume of 100 μl per dose and loaded into syringes with a 26-gauge needle for subcutaneous injection. Mice were boosted on Week 3. For hydrogel groups, vaccines were prepared as described above to a volume of 100 μl per dose (except for the double volume group, where two doses of antigen and a dose of adjuvant were prepared to a volume of 200 μL) in syringes with a 21-gauge needle for subcutaneous injection.

### 5.9 Mice and Vaccination

Six-to-seven weeks old female C57BL/6 (B6) mice were purchased from Charles River and housed in the animal facility at Stanford University. Mice were shaved to receive a subcutaneous injection of 100 µL of soluble or hydrogel vaccine on the right side of their flank under brief anesthesia. Mouse blood was collected weekly from the tail veins.

### 5.10 Mouse Serum ELISAs

Serum antigen-specific IgG antibody endpoint titers were measured using an endpoint ELISA. MaxiSorp plates (Invitrogen) were coated with SARS-CoV-2 RBD protein, spike protein, or a spike protein variant (B.1.617.2, or B.1.529), at 2 µg/mL in PBS 1X overnight at 4 °C and subsequently blocked with PBS 1X containing 1 wt% BSA for 1 h at 25 °C. Plates were washed 5 times in between each step with PBS 1X containing 0.05 wt% Tween-20. Serum samples were diluted in diluent buffers (PBS 1X with 1 wt% BSA) starting at 1:100 and 4-fold serially diluted and incubated in the previously coated plates for 2 h at 25 °C. Goat-anti-mouse IgG Fc-HRP (1:10,000), IgG1 Fc-HRP (1:10,000), or IgG2c-HRP (1:10,000) were then added for 1 h at 25 °C. The plates were developed with TMB substrate, and the reaction stopped with 1 M HCl. The absorbances were analyzed using a Synergy H1 microplate reader (BioTek Instruments) at 450 nm. The total IgG, the subtypes, and the variants were imported into GraphPad Prism 8.4.1 to determine the serum titers by fitting the curves with a three-parameter non-linear regression (baseline constrained to 0.054, the negative control average). The dilution titer value at which the endpoint threshold (0.1) was crossed for each curve was imputed. Samples failing to meet endpoint threshold at a 1:100 dilution were set to a titer cutoff of 1:25 or below the limit quantitation for the assay.

### 5.11 Antibody Half-life Calculations

The Power law decay model was used to estimate the decay half-life of binding antibody endpoint titer. The equation *d*Ab/*d*t = -*k*/*t* × Ab and Ab = C × *t*^-*k*^ were fitted using Matlab’s fitnlm function (R2022b, Mathworks) to the longitudinal data starting from D56 after the initial immunization (for which we observed antibody endpoint titers to be at their peak). Ab is the RBD-specific antibody binding endpoint titers and *k* is the power law decay rate. Longitudinal data from three time points, D56, D70, and D182 (6M) were used to estimate the decay rate. The corresponding half-lives were calculated as *t*_1/2_=0.5^1/*k*^.

### 5.12 SARS-CoV-2 Spike-Pseudotyped Viral Neutralization

SARS-CoV-2 spike-pseudotyped lentivirus production and neutralization assays were performed as described previously.[69] Briefly, SARS-CoV-2 spike-pseudotyped lentivirus was produced in HEK293T cells using a five-plasmid system.[31] After plating 6 million HEK293T cells overnight, the cells were transfected with DNA using BioT (BioLand). Virus-containing supernatants were collected and filtered through a 0.45 µm filter 72 h post-transfection. The virus was aliquoted and stored at −80 °C.

Neutralization assays were done using HeLa cells expressing ACE2 and TMPRSS2. Cells were plated in a 96-well plate at a density of 8000 cells/well 1 day prior to infection. Serum was heat inactivated for 15 min at 56 °C before dilution in cell culture medium and then 60 µL of heat-inactivated serum was mixed with 60 µL of virus and polybrene. Polybrene (Sigma-Aldrich, Cat # TR-1003-G) was present at a final concentration of 5 µg/mL in all samples. Serum/virus dilutions were incubated at 37 °C for 1 h before 100 µL from each well was added to a 96-well plate seeded with cells. After 2 days at 37 °C, cells were lysed using BriteLite (Perkin Elmer) reagent and luminescence was measured using a BioTek Synergy HT Microplate Reader (BioTek). Each plate was normalized by averaging cell-only wells (0 % infectivity) and virus-only wells (100 % infectivity). Normalized values were plotted and fitted in Prism with a three-parameter non-linear regression curve to obtain 50 % inhibitor concentration (NT_50_) values.

### 5.13 Animal Protocol

Mice were cared for according to Institutional Animal Care and Use guidelines. All animal studies were performed in accordance with the National Institutes of Health guidelines and the approval of the Stanford Administrative Panel on Laboratory Animal Care (Protocol APLAC-32109).

### 5.14 Collection of Human Convalescent Serum for Previously Infected Human Patients

Convalescent COVID-19 blood was collected from donors 8–12 weeks after the onset of symptoms. Blood was collected in microtubes with serum gel for clotting (Starstedt), centrifuged for 5 mins at 10,000 g and then stored at −80 °C until used. Blood collection was done by finger-prick and was performed in accordance with National Institutes of Health guidelines with the approval of the Stanford Human Subjects Research and IRB Compliance Office (IRB-58511) and with the consent of the individuals.

### 5.15 Statistical Analysis

For *in vivo* experiments, animals were cage blocked. All results are expressed as mean ± standard error of mean (SEM). Comparisons between two groups were conducted by a two-tailed Student’s t-test. One-way ANOVA tests with a Tukey’s multiple-comparisons test were used for comparison across multiple groups. For plots displaying multiple time points or protection against different variants, *p* values were determined with a 2way ANOVA with Tukey’s multiple-comparisons test. Statistical analysis was performed using GraphPad Prism 8.4.1 (GraphPad Software). Statistical significance was considered as *p* < 0.05. Selected *p* values are shown in the text and reported in the supporting information.

## Data Availability

The data that support the findings of this study are available from the corresponding author upon reasonable request.

## Supporting Information

The following files are available free of charge. Additional tables and figures of synthesis, purification, and characterization of PNP hydrogels and additional immune response data and figures. (PDF)

## Author Contributions

BO and EA designed the broad concepts and research studies. OS, JY, and TB designed specific experiments. BO, OS, JY, and TB performed research and experiments. BO and EA wrote the paper. OS, JY, and JB edited the paper. The manuscript was written through contributions of all authors. All authors have given approval to the final version of the manuscript.

## Acknowledgment

We would like to thank all members of the Appel lab for their useful discussion and advice throughout this project. Also, the staff of the BioE/ChemE Animal Facility who cared for our mice. The authors would also like to thank the Stanford Core Facilities that were central to the completion of this work. We also would like to thank the human patients for donating their serums. TB thanks Ya-Chen Cheng for making SARS-CoV-2 WT lentivirus. We would also like to thank Jeremy Blum for his countless advice and Christopher Fox for both access to 3M-052 and his assistances. We are also thankful for Professor Peter Kim for his assistance in conducting the neutralization studies in the work. This work was financially supported by the Center for Human Systems Immunology with the Bill & Melinda Gates Foundation (OPP1113682; OPP1211043; INV027411), NIH shared instrumentation grant (S10OD021600), and a Bio-X Interdisciplinary Initiatives Seed Grant. BO is grateful for an Eastman Kodak Fellowship. OS and JY are thankful for a National Science Foundation Graduate Research Fellowship. TB is thankful for the Knight-Hennessy Graduate Scholarship and a Canadian Institutes of Health Research Doctoral Foreign Study Award (FRN:170770). JB is thankful for a Marie-Curie fellowship from the European Union (H2020; No. 101030481).

