## Supplemental Information for "Broad and Durable Humoral Responses Following Single Hydrogel Immunization of SARS-CoV-2 Subunit Vaccine"

**Supplemental Figures**

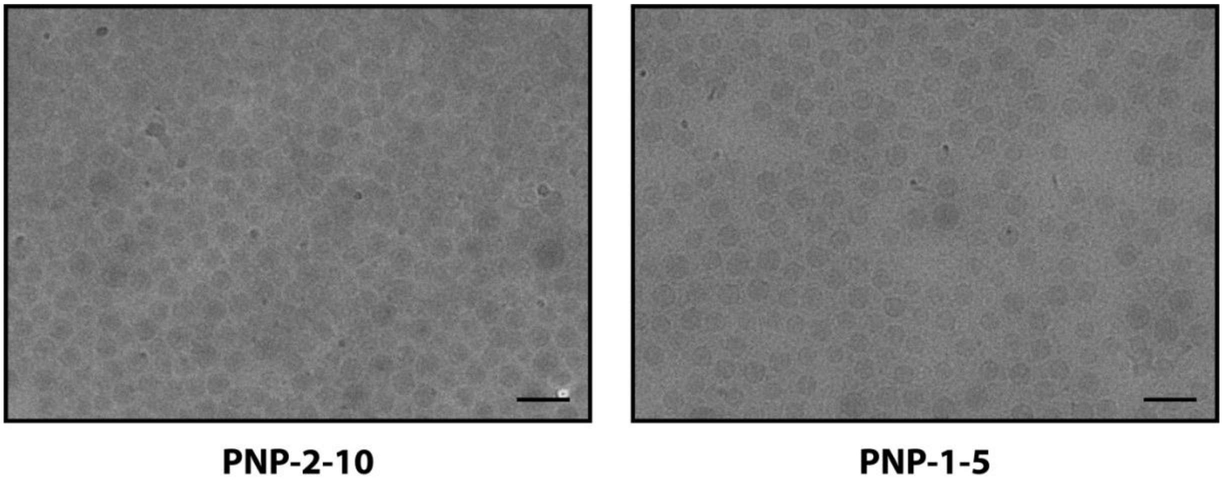


**Figure S1. CryoEM images of PNP hydrogels.** CryoEM images showing nanoparticles are evenly dispersed in the polymer matrix and not agglomerated. Scale bar: 50nm**.**

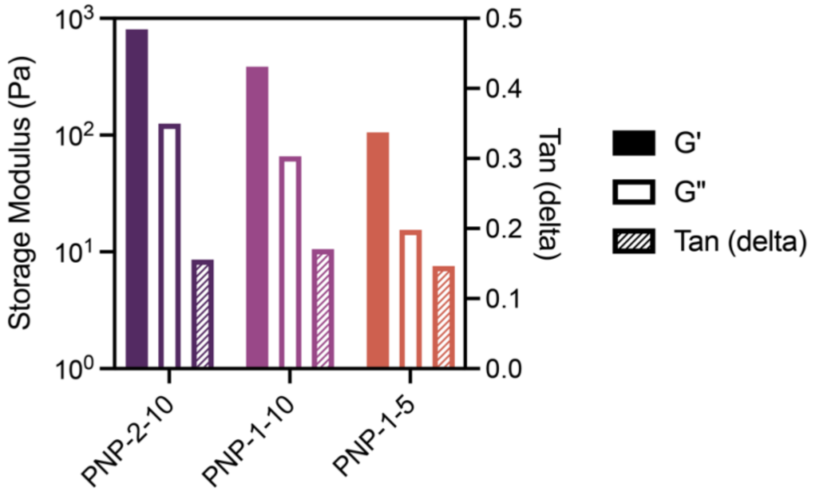


**Figure S2. PNP hydrogels exhibit solid-like properties.** Storage (G'), Loss (G"), and Tan (delta) of PNP hydrogel formulations at an angular frequency of $\omega$ = 10 rad/s**.**

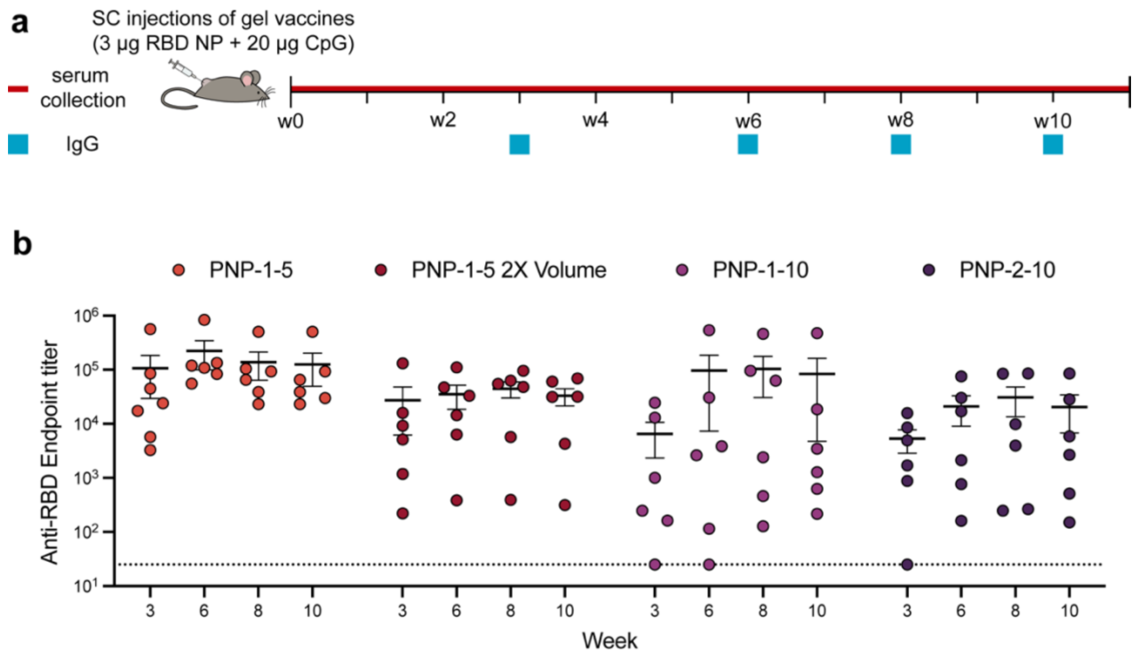


**Figure S3. PNP hydrogel vaccines *in vivo* humoral responses with RBD NP. (a)** Expended timeline of mouse immunizations and blood collection. Mice were immunized with PNP hydrogels formulated with 3 µg of RBD NP on day 0 and serum was collected over time. **(b)** Anti-RBD IgG binding endpoint titers of different PNP hydrogel formulations, all with 20 µg of CpG. Data are shown as mean +/- SEM. *p* values were determined using a 2way ANOVA with Tukey's multiple comparisons test on the logged titer values for IgG titer comparisons. *p* values for comparisons are shown in Table S3.

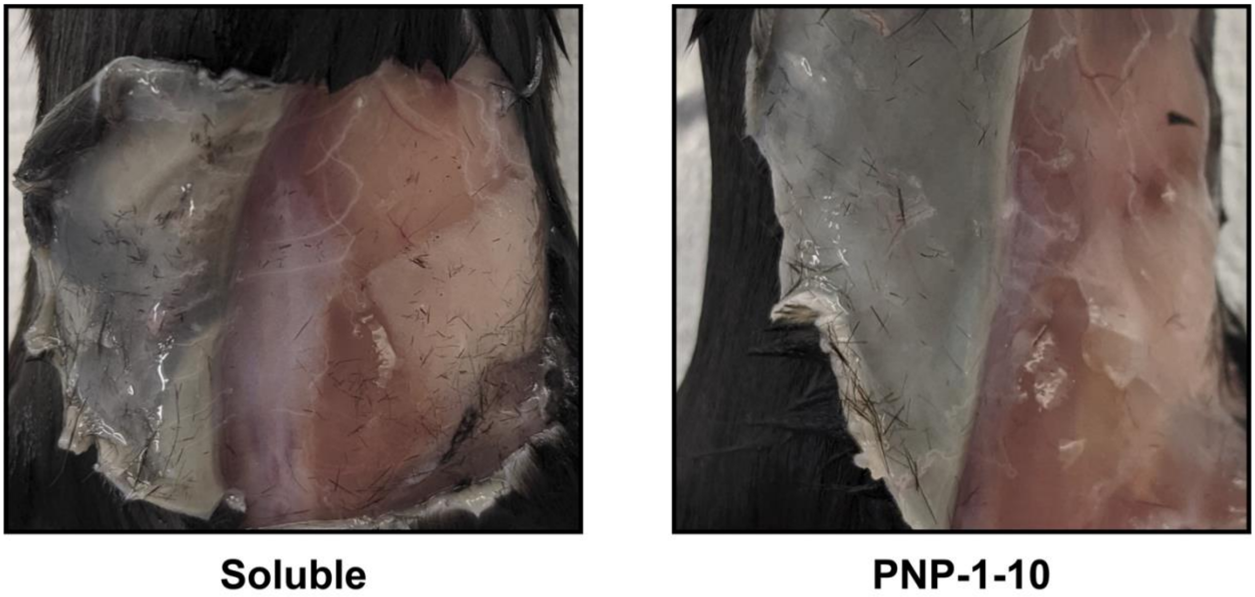


**Figure S4. Long-term biocompatibility in subcataneous space.** Image of the subcutaneous space 12 weeks after adminstration of either saline or PNP-1-10 Hydrogel. Images indicate no noticeable vascularization or fibrotic responses following hydrogel treatments, and there were no differences between hydrogel and control treatments. The hydrogel materials were completely degraded by this time point. Similar images demonstrating biocomptability of PNP-2-10 and PNP-1-5 have been previously reported.[1]

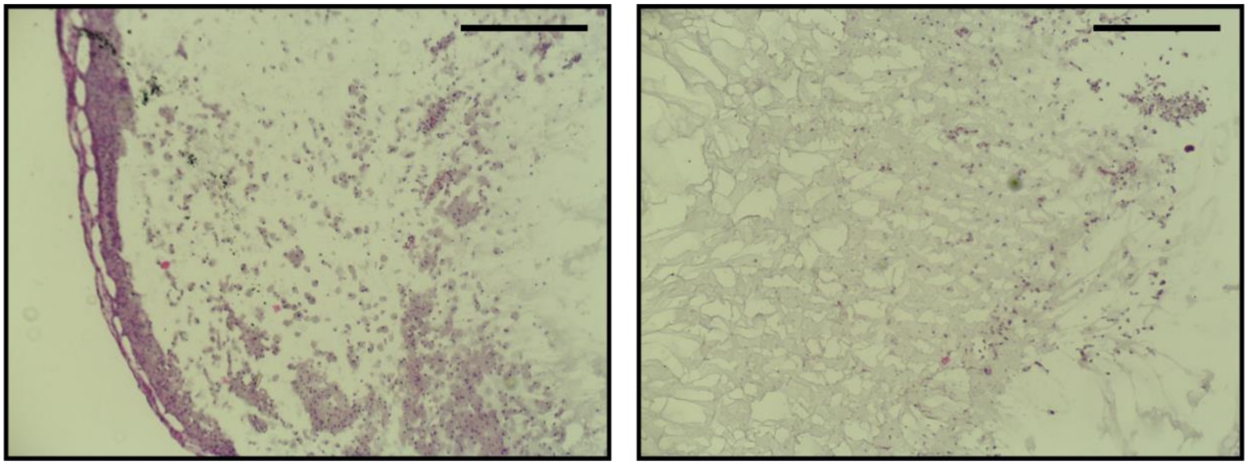

**Figure S5. Histology images of PNP-2-10.** PNP-2-10 hydrogels were explanted from the subcutaneous space 7 days after immunization. Images demonstrate that no fibrosis or vasculature has formed around the hydrogel, indicating excellent biocompatibility of these materials. Additionally, cells have begun to infiltrate the gel, with a greater concentration at the outer edge. Additional histology images demonstrating biocompatibility of PNP hydrogels have been previously reported.[2] Scale bars: 400nm.

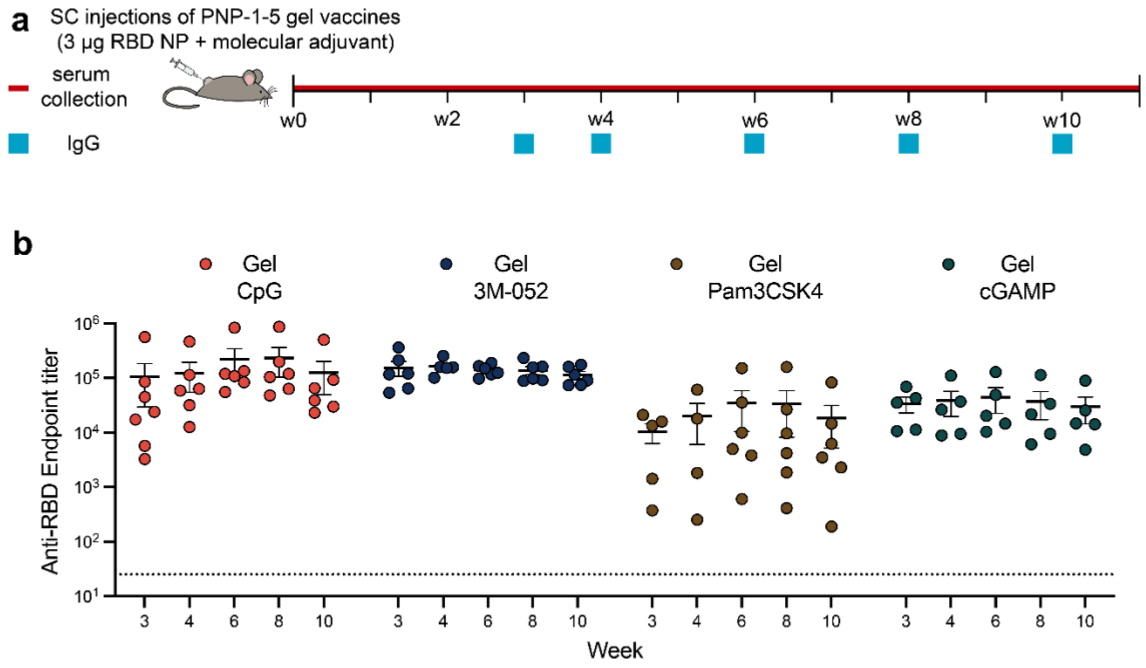


**Figure S6. PNP hydrogel vaccines *in vivo* humoral responses with RBD NP. (a)** Expended timeline of mouse immunizations and blood collection. Mice were immunized with PNP hydrogels formulated with 3 μg of RBD NP on day 0 and serum was collected over time. **(b)** Anti-RBD IgG binding endpoint titers of PNP-1-5 hydrogels formulated with different clinical relevant molecular adjuvants. Data are shown as mean +/- SEM. *p* values were determined using a 2way ANOVA with Tukey's multiple comparisons test on the logged titer values for IgG titer comparisons. *p* values for comparisons are shown in Table S4.

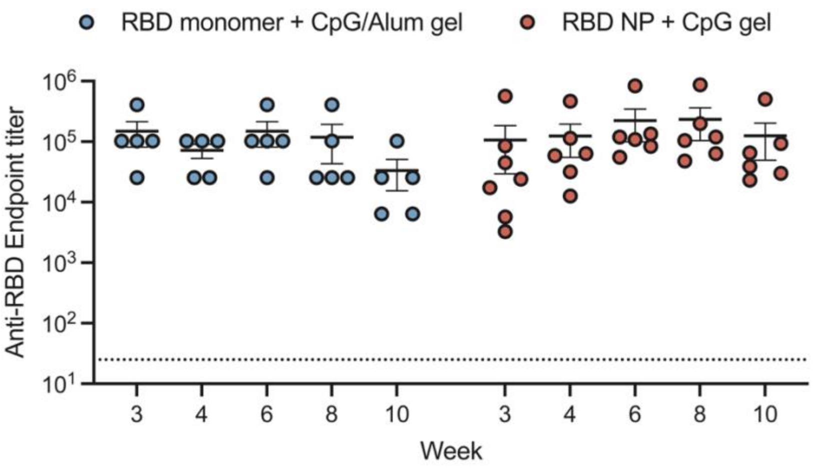


**Figure S7. RBD NP hydrogel vaccine is more potent and durable than RBD monomer hydrogel vaccine.** Anti-RBD IgG binding endpoint titers of previously reported[3] RBD monomer hydrogel and RBD NP hydrogel. Data are shown as mean +/- SEM. *p* values were determined using a 2way ANOVA with Tukey's multiple comparisons test on the logged titer values for IgG titer comparisons. *p* values for comparisons are shown in Tables S5 & S6.

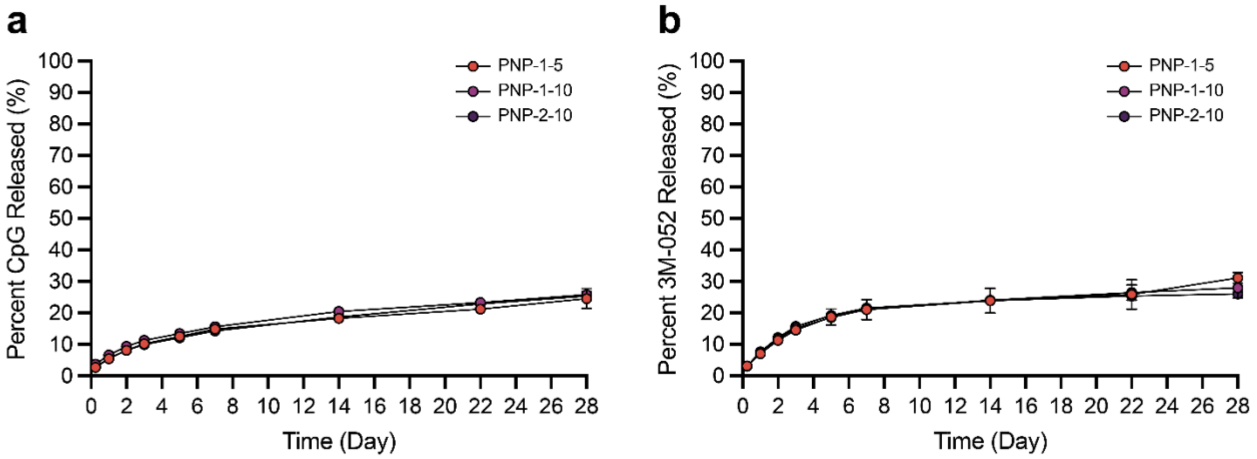


**Figure S8. PNP Hydrogels limit adjuvants diffusion.** Release kinetics of **(a)** CpG and **(b)** 3M-052 from PNP hydrogels in a glass capillary *in vitro* release study.

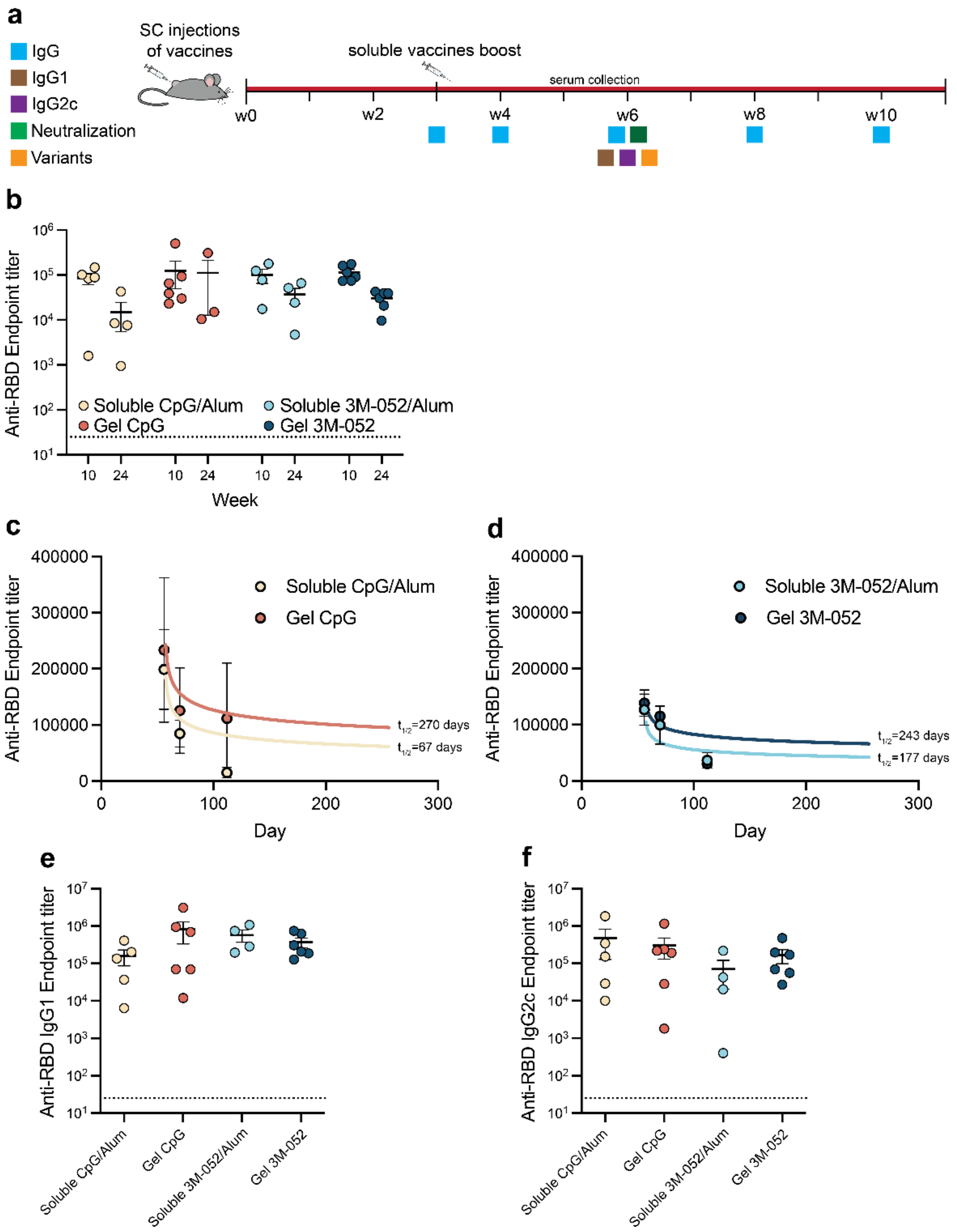


**Figure S9. Single immunization PNP hydrogel vaccine is durable over 6 months period. (a)** Timeline of mouse immunizations and blood collection to determine IgG titers. Mice were immunized with either PNP-1-5 hydrogels formulated with 3 μg of RBD NP on day 0 or soluble vaccines formulated with 1.5 μg of RBD on day 0 and day 21. IgG1, IgG2c, neutralization, and variants titers were determined on day 42. **(b)** Anti-RBD IgG binding endpoint titers of treatment groups on Week 10 and Week 24 (6 months). **(c-d)** The power law decay model was used to determine the binding antibody decay half-lives for the treatment groups. Week 8, Week 10, and Week 24 time points were used for the fit. **(e)** Anti-RBD IgG1 and **(d)** IgG2c titers from serum collected on week 6 after immunization. Data are shown as mean +/- SEM. *p* values were determined using a 2way ANOVA with Tukey's multiple comparisons test on the logged titer values for IgG titer comparisons. *p* values for comparisons are shown in Tables S8-S12.

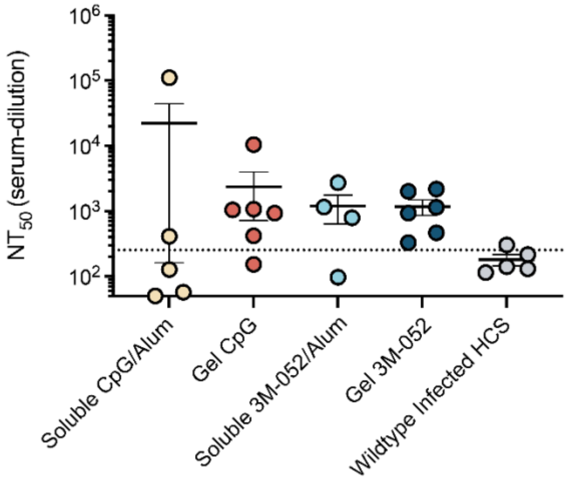


**Figure S10. Single immunization hydrogel vaccines elicit robust neutralizing antibodies in mice.** Comparison of $NT_{50}$ values determined from neutralization curves. Human Convalescent sera $NT_{50}$ values were previously reported.[3] Data are shown as mean +/- SEM. *p* values were determined using a 2way ANOVA with Tukey's multiple comparisons test on the logged titer values for IgG titer comparisons. *p* values for comparisons are shown in Table S14.

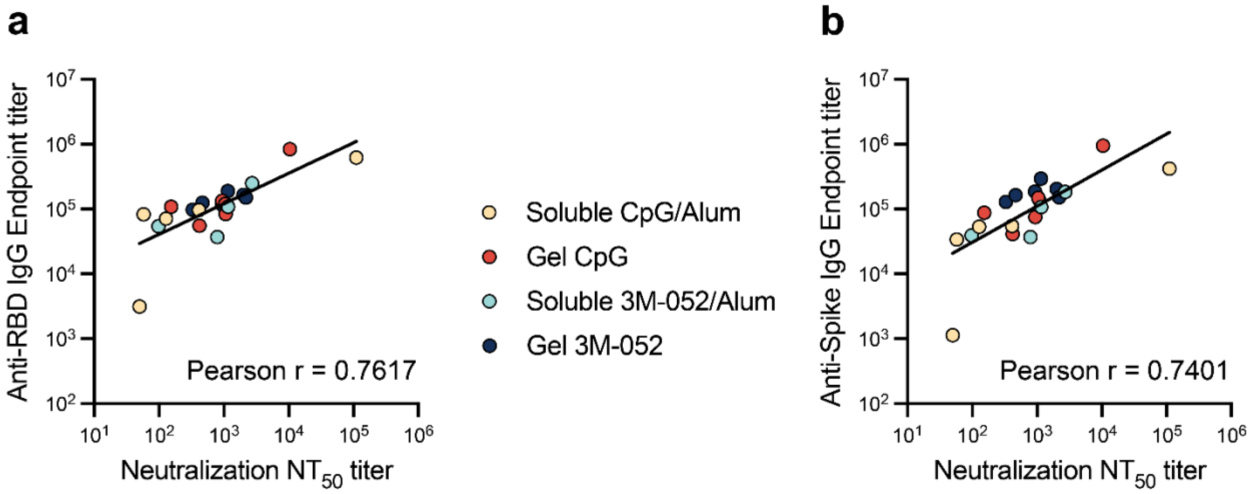


**Figure 11. RBD and spike binding titers are highly correlative with neutralization $NT_{50}$ titers.** Week 6 **(a)** anti-RBD and **(b)** anti-spike binding titers plotted against neutralization $NT_{50}$ titer and Pearson correlations were determined.

### Supplemental Tables

### Table S1- Vaccine groups

| Vaccine Group Name | Number of admin | Vehicle | Hydrogel formulation (% HPMC-$C_{12}$ - % PEG-*b*-PLA) | Antigen Dose (µg) | Adjuvant | Adjuvant dose (µg) |
|---|---|---|---|---|---|---|
| **PNP-1-5 / Gel CpG** | 1 | Hydrogel | 1-5 | 3 | CpG1826 | 20 |
| **PNP-1-5 2X Volume** | 1 | Hydrogel | 1-5 | 3 | CpG1826 | 20 |
| **PNP-1-10** | 1 | Hydrogel | 1-10 | 3 | CpG1826 | 20 |
| **PNP-2-10** | 1 | Hydrogel | 2-10 | 3 | CpG1826 | 20 |
| **Gel Pam3CSK4** | 1 | Hydrogel | 1-5 | 3 | Pam3CSK4 | 20 |
| **Gel 3M-052** | 1 | Hydrogel | 1-5 | 3 | 3M-052 | 1 |
| **Gel cGAMP** | 1 | Hydrogel | 1-5 | 3 | cGAMP | 20 |
| **Soluble CpG/Alum** | 2 | Alum | - | 1.5 x 2 | CpG1826/ Alum | 20/100 |
| **Soluble 3M-052/Alum** | 2 | Alum | - | 1.5 x 2 | 3M-052/Alum | 1/100 |

**Table S2**- DLS measurements of 500,000MW FITC-Dextran.

| Diameter (nm) | Polydispersity (nm) |
|---|---|
| 39.4 | 5.5 |
| 40.6 | 7.3 |
| 38.4 | 4.3 |
| 39.2 | 6.4 |
| 38.3 | 4.3 |

Average ± SEM: 39.2 ± 0.4

**Table S3**- *p* values from a 2-way ANOVA with Tukey's multiple comparisons test for specific IgG titer time points compared between among PNP hydrogel formulations (referring to Figures 3b and S3).

| Week 3<br>IgG titers [$\log_{10}$] | Adjusted *p* values |
|---|---|
| PNP-1-5 vs. PNP-1-5 2X Volume | 0.5667 |
| PNP-1-5 vs. PNP-1-10 | 0.0351 |
| PNP-1-5 vs. PNP-2-10 | 0.1324 |
| PNP-1-5 2X Volume vs. PNP-1-10 | 0.4967 |
| PNP-1-5 2X Volume vs. PNP-2-10 | 0.8217 |
| PNP-1-10 vs. PNP-2-10 | 0.9482 |

| Week 6<br>IgG titers [$\log_{10}$] | Adjusted *p* values |
|---|---|
| PNP-1-5 vs. PNP-1-5 2X Volume | 0.3241 |
| PNP-1-5 vs. PNP-1-10 | 0.0275 |
| PNP-1-5 vs. PNP-2-10 | 0.0699 |
| PNP-1-5 2X Volume vs. PNP-1-10 | 0.6528 |
| PNP-1-5 2X Volume vs. PNP-2-10 | 0.8611 |
| PNP-1-10 vs. PNP-2-10 | 0.9815 |

| Week 8<br>IgG titers [$\log_{10}$] | Adjusted *p* values |
|---|---|
| PNP-1-5 vs. PNP-1-5 2X Volume | 0.6857 |
| PNP-1-5 vs. PNP-1-10 | 0.3516 |
| PNP-1-5 vs. PNP-2-10 | 0.1849 |
| PNP-1-5 2X Volume vs. PNP-1-10 | 0.9448 |
| PNP-1-5 2X Volume vs. PNP-2-10 | 0.7880 |
| PNP-1-10 vs. PNP-2-10 | 0.9816 |

| Week 10<br>IgG titers [$\log_{10}$] | Adjusted *p* values |
|---|---|
| PNP-1-5 vs. PNP-1-5 2X Volume | 0.6403 |
| PNP-1-5 vs. PNP-1-10 | 0.1850 |
| PNP-1-5 vs. PNP-2-10 | 0.1601 |
| PNP-1-5 2X Volume vs. PNP-1-10 | 0.8264 |
| PNP-1-5 2X Volume vs. PNP-2-10 | 0.7875 |
| PNP-1-10 vs. PNP-2-10 | 0.9998 |

**Table S4**- *p* values from a 2-way ANOVA with Tukey's multiple comparisons test for specific IgG titer time points compared among different PNP hydrogel adjuvant formulations (referring to Figures 3c and S6).

| Week 3<br>IgG titers [$\log_{10}$] | Adjusted *p* values |
|---|---|
| Gel CpG vs. Gel 3M-052 | 0.1941 |
| Gel CpG vs. Gel Pam3CSK4 | 0.0922 |
| Gel CpG vs. Gel cGAMP | 0.9993 |
| Gel 3M-052 vs. Gel Pam3CSK4 | 0.0005 |
| Gel 3M-052 vs. Gel cGAMP | 0.2091 |
| Gel Pam3CSK4 vs. Gel cGAMP | 0.1717 |

| Week 4<br>IgG titers [$\log_{10}$] | Adjusted *p* values |
|---|---|
| Gel CpG vs. Gel 3M-052 | 0.6849 |
| Gel CpG vs. Gel Pam3CSK4 | 0.0133 |
| Gel CpG vs. Gel cGAMP | 0.6044 |
| Gel 3M-052 vs. Gel Pam3CSK4 | 0.0008 |
| Gel 3M-052 vs. Gel cGAMP | 0.1183 |
| Gel Pam3CSK4 vs. Gel cGAMP | 0.2480 |

| Week 6<br>IgG titers [$\log_{10}$] | Adjusted *p* values |
|---|---|
| Gel CpG vs. Gel 3M-052 | >0.9999 |
| Gel CpG vs. Gel Pam3CSK4 | 0.0029 |
| Gel CpG vs. Gel cGAMP | 0.1874 |
| Gel 3M-052 vs. Gel Pam3CSK4 | 0.0028 |
| Gel 3M-052 vs. Gel cGAMP | 0.1866 |
| Gel Pam3CSK4 vs. Gel cGAMP | 0.4615 |

| Week 8<br>IgG titers [$\log_{10}$] | Adjusted *p* values |
|---|---|
| Gel CpG vs. Gel 3M-052 | 0.9999 |
| Gel CpG vs. Gel Pam3CSK4 | 0.0010 |
| Gel CpG vs. Gel cGAMP | 0.0909 |
| Gel 3M-052 vs. Gel Pam3CSK4 | 0.0012 |
| Gel 3M-052 vs. Gel cGAMP | 0.1055 |
| Gel Pam3CSK4 vs. Gel cGAMP | 0.4753 |

| Week 10<br>IgG titers [$\log_{10}$] | Adjusted *p* values |
|---|---|
| Gel CpG vs. Gel 3M-052 | 0.9057 |
| Gel CpG vs. Gel Pam3CSK4 | 0.0038 |
| Gel CpG vs. Gel cGAMP | 0.3673 |
| Gel 3M-052 vs. Gel Pam3CSK4 | 0.0004 |
| Gel 3M-052 vs. Gel cGAMP | 0.1148 |
| Gel Pam3CSK4 vs. Gel cGAMP | 0.2923 |

**Table S5-** *p* values from a multiple unpaired Welch's-t test for specific IgG titer time points compared between the RBD Monomer PNP hydrogel and the RBD NP PNP hydrogel groups (referring to Figure S7).

| RBD monomer + CpG/Alum gel vs. RBD NP + CpG gel IgG titers [$\log_{10}$] | Adjusted *p* values |
|---|---|
| Week 3 | 0.1402 |
| Week 4 | 0.8599 |
| Week 6 | 0.6266 |
| Week 8 | 0.2620 |
| Week 10 | 0.1119 |

**Table S6**- *p* values from a 2-way ANOVA with Dunnett's multiple comparisons test for the RBD Monomer PNP hydrogel and the RBD NP PNP hydrogel groups' specific IgG titers for which Week 3, Week 4, Week 6, and Week 8 titers are compared to Week 10 titers (referring to Figure S7).

| RBD monomer + CpG/Alum gel<br>IgG titers [$\log_{10}$] | Adjusted *p* values |
|---|---|
| Week 10 vs. Week 3 | 0.0904 |
| Week 10 vs. Week 4 | 0.3710 |
| Week 10 vs. Week 6 | 0.0904 |
| Week 10 vs. Week 8 | 0.3710 |

| Week 10<br>IgG titers [$\log_{10}$] | Adjusted *p* values |
|---|---|
| Week 10 vs. Week 3 | 0.5240 |
| Week 10 vs. Week 4 | >0.9999 |
| Week 10 vs. Week 6 | 0.6563 |
| Week 10 vs. Week 8 | 0.6631 |

**Table S7**- *p* values from a 2-way ANOVA with Tukey's multiple comparisons test for specific IgG titer time points compared between different vaccines (referring to Figure 4b).

| Week 1<br>IgG titers [$log_{10}$] | Adjusted *p* values |
|---|---|
| Soluble CpG/Alum vs. Soluble 3M-052/Alum | 0.9667 |
| Soluble CpG/Alum vs. Gel CpG | 0.9618 |
| Soluble CpG/Alum vs. Gel 3M-052 | 0.2914 |
| Soluble 3M-052/Alum vs. Gel CpG | >0.9999 |
| Soluble 3M-052/Alum vs. Gel 3M-052 | 0.5724 |
| Gel CpG vs. Gel 3M-052 | 0.4569 |

| Week 2<br>IgG titers [$log_{10}$] | Adjusted *p* values |
|---|---|
| Soluble CpG/Alum vs. Soluble 3M-052/Alum | 0.3371 |
| Soluble CpG/Alum vs. Gel CpG | 0.0064 |
| Soluble CpG/Alum vs. Gel 3M-052 | <0.0001 |
| Soluble 3M-052/Alum vs. Gel CpG | 0.4692 |
| Soluble 3M-052/Alum vs. Gel 3M-052 | 0.0131 |
| Gel CpG vs. Gel 3M-052 | 0.2217 |

| Week 3<br>IgG titers [$log_{10}$] | Adjusted *p* values |
|---|---|
| Soluble CpG/Alum vs. Soluble 3M-052/Alum | 0.2096 |
| Soluble CpG/Alum vs. Gel CpG | <0.0001 |
| Soluble CpG/Alum vs. Gel 3M-052 | <0.0001 |
| Soluble 3M-052/Alum vs. Gel CpG | 0.0155 |
| Soluble 3M-052/Alum vs. Gel 3M-052 | 0.0002 |
| Gel CpG vs. Gel 3M-052 | 0.3765 |

| Week 4<br>IgG titers [$log_{10}$] | Adjusted *p* values |
|---|---|
| Soluble CpG/Alum vs. Soluble 3M-052/Alum | 0.5434 |
| Soluble CpG/Alum vs. Gel CpG | 0.1060 |
| Soluble CpG/Alum vs. Gel 3M-052 | 0.0155 |
| Soluble 3M-052/Alum vs. Gel CpG | 0.8675 |
| Soluble 3M-052/Alum vs. Gel 3M-052 | 0.4139 |
| Gel CpG vs. Gel 3M-052 | 0.8084 |

| Week 6<br>IgG titers [$log_{10}$] | Adjusted *p* values |
|---|---|
| Soluble CpG/Alum vs. Soluble 3M-052/Alum | 0.9937 |
| Soluble CpG/Alum vs. Gel CpG | 0.8723 |
| Soluble CpG/Alum vs. Gel 3M-052 | 0.8715 |
| Soluble 3M-052/Alum vs. Gel CpG | 0.9711 |
| Soluble 3M-052/Alum vs. Gel 3M-052 | 0.9708 |
| Gel CpG vs. Gel 3M-052 | >0.9999 |

| Week 8<br>IgG titers [$log_{10}$] | Adjusted *p* values |
|---|---|
| Soluble CpG/Alum vs. Soluble 3M-052/Alum | >0.9999 |
| Soluble CpG/Alum vs. Gel CpG | 0.9964 |
| Soluble CpG/Alum vs. Gel 3M-052 | 0.9984 |
| Soluble 3M-052/Alum vs. Gel CpG | 0.9985 |
| Soluble 3M-052/Alum vs. Gel 3M-052 | 0.9995 |
| Gel CpG vs. Gel 3M-052 | >0.9999 |

| Week 10<br>IgG titers [$log_{10}$] | Adjusted *p* values |
|---|---|
| Soluble CpG/Alum vs. Soluble 3M-052/Alum | 0.9672 |
| Soluble CpG/Alum vs. Gel CpG | 0.9792 |
| Soluble CpG/Alum vs. Gel 3M-052 | 0.8089 |
| Soluble 3M-052/Alum vs. Gel CpG | 0.9995 |
| Soluble 3M-052/Alum vs. Gel 3M-052 | 0.9842 |
| Gel CpG vs. Gel 3M-052 | 0.9532 |

**Table S8**- *p* values from a Welch's-t test for Week 24 IgG titer compared between the Soluble and the Gel groups (referring to Figure S9b).

| Week 24<br>IgG titers [$log_{10}$] | Adjusted *p* values |
|---|---|
| Soluble CpG/Alum vs. Gel CpG | 0.2878 |
| Soluble 3M-052/Alum vs. Gel 3M-052 | 0.8924 |

**Table S9**- *p* values from a Welch's-t test for the antibody decay half-life compared between the Soluble and the Gel groups (referring to Figures 4c, S9b, S9c, and S9d).

| Antibody decay half-life | Adjusted *p* values |
|---|---|
| Soluble CpG/Alum vs. Gel CpG | 0.3905 |
| Soluble 3M-052/Alum vs. Gel 3M-052 | 0.7662 |

**Table S10**- *p* values from a 1-way ANOVA with Tukey's multiple comparisons test for specific IgG1 titer on week 6 compared between different soluble and hydrogel vaccines (referring to Figure S9e).

| Week 6<br>IgG1 titers [$log_{10}$] | Adjusted *p* values |
|---|---|
| Soluble CpG/Alum vs. Soluble 3M-052/Alum | 0.3009 |
| Soluble CpG/Alum vs. Gel CpG | 0.6374 |
| Soluble CpG/Alum vs. Gel 3M-052 | 0.4428 |
| Soluble 3M-052/Alum vs. Gel CpG | 0.8697 |
| Soluble 3M-052/Alum vs. Gel 3M-052 | 0.9686 |
| Gel CpG vs. Gel 3M-052 | 0.9848 |

**Table S11**- *p* values from a 1-way ANOVA with Tukey's multiple comparisons test for specific IgG2c titer on week 6 compared between different soluble and hydrogel vaccines (referring to Figure S9f).

| Week 6<br>IgG2c titers [$\log_{10}$] | Adjusted *p* values |
|---|---|
| Soluble CpG/Alum vs. Soluble 3M-052/Alum | 0.4689 |
| Soluble CpG/Alum vs. Gel CpG | 0.9950 |
| Soluble CpG/Alum vs. Gel 3M-052 | 0.9996 |
| Soluble 3M-052/Alum vs. Gel CpG | 0.5656 |
| Soluble 3M-052/Alum vs. Gel 3M-052 | 0.4891 |
| Gel CpG vs. Gel 3M-052 | 0.9988 |

**Table S12**- *p* values from a 1-way ANOVA with Tukey's multiple comparisons test for the ratio of IgG2c/IgG1 titer on week 6 compared between different soluble and hydrogel vaccines (referring to Figure 4d).

| Week 6<br>IgG2c/IgG1 ratio [$log_{10}$] | Adjusted *p* values |
|---|---|
| Soluble CpG/Alum vs. Soluble 3M-052/Alum | 0.1063 |
| Soluble CpG/Alum vs. Gel CpG | 0.7689 |
| Soluble CpG/Alum vs. Gel 3M-052 | 0.7184 |
| Soluble 3M-052/Alum vs. Gel CpG | 0.3886 |
| Soluble 3M-052/Alum vs. Gel 3M-052 | 0.4335 |
| Gel CpG vs. Gel 3M-052 | 0.9997 |

**Table S13-** *p* values from a 2-way ANOVA with Tukey's multiple comparisons test for specific spike variant titers compared between different soluble and hydrogel vaccines (referring to Figure 5).

| WT Spike IgG titers [$\log_{10}$] | Adjusted *p* values |
|---|---|
| Soluble CpG/Alum vs. Soluble 3M-052/Alum | 0.8405 |
| Soluble CpG/Alum vs. Gel CpG | 0.3677 |
| Soluble CpG/Alum vs. Gel 3M-052 | 0.1969 |
| Soluble 3M-052/Alum vs. Gel CpG | 0.9065 |
| Soluble 3M-052/Alum vs. Gel 3M-052 | 0.7348 |
| Gel CpG vs. Gel 3M-052 | 0.9790 |

| Delta (B.617.2) IgG titers [$\log_{10}$] | Adjusted *p* values |
|---|---|
| Soluble CpG/Alum vs. Soluble 3M-052/Alum | 0.7375 |
| Soluble CpG/Alum vs. Gel CpG | 0.5432 |
| Soluble CpG/Alum vs. Gel 3M-052 | 0.4321 |
| Soluble 3M-052/Alum vs. Gel CpG | 0.1074 |
| Soluble 3M-052/Alum vs. Gel 3M-052 | 0.0736 |
| Gel CpG vs. Gel 3M-052 | 0.9974 |

| Omicron (B.1.1.529) IgG titers [$\log_{10}$] | Adjusted *p* values |
|---|---|
| Soluble CpG/Alum vs. Soluble 3M-052/Alum | 0.3526 |
| Soluble CpG/Alum vs. Gel CpG | 0.2173 |
| Soluble CpG/Alum vs. Gel 3M-052 | 0.0311 |
| Soluble 3M-052/Alum vs. Gel CpG | 0.9996 |
| Soluble 3M-052/Alum vs. Gel 3M-052 | 0.7822 |
| Gel CpG vs. Gel 3M-052 | 0.7867 |

**Table S14**- *p* values from a 1-way ANOVA with Tukey's multiple comparisons test for specific $NT_{50}$ neutralization titer on week 6 compared between different soluble and hydrogel vaccines (referring to Figure 6e and Figure S10).

| Week 6<br>$NT_{50}$ titers [$\log_{10}$] | Adjusted *p* values |
|---|---|
| Soluble CpG/Alum vs. Soluble 3M-052/Alum | 0.9937 |
| Soluble CpG/Alum vs. Gel CpG | 0.9445 |
| Soluble CpG/Alum vs. Gel 3M-052 | 0.9435 |
| Soluble 3M-052/Alum vs. Gel CpG | 0.9987 |
| Soluble 3M-052/Alum vs. Gel 3M-052 | 0.9987 |
| Gel CpG vs. Gel 3M-052 | >0.9999 |
| Wildtype Infected HCS vs. Soluble CpG/Alum | 0.8955 |
| Wildtype Infected HCS vs. Soluble 3M-052/Alum | 0.7215 |
| Wildtype Infected HCS vs. Gel CpG | 0.4695 |
| Wildtype Infected HCS vs. Gel 3M-052 | 0.4673 |

### Supplemental Methods

#### Histology of PNP-2-10 Hydrogels

A cohort of five 8-10-week-old C57BL/6 mice were inoculated subcutaneously with 100 μL of PNP-2-10 hydrogels. Hydrogels were explanted from the subcutaneous space 7 days after injection and immediately frozen in optimal cutting temperature compound. Samples were sectioned, mounted on glass slides, and stained with H&E staining. Images were taken using a 10x objective on an optical microscope.
